# FXR proteins and BICD2 cooperate to promote dynein-mediated RNP transport and stress granule assembly

**DOI:** 10.64898/2026.09.25.754448

**Authors:** Yaiza Andrés Jeske, Eve Mehtab, Lucas Albacete-Albacete, Mark A. McClintock, Jérôme Boulanger, Simon L. Bullock

## Abstract

Intracellular trafficking of ribonucleoprotein particles (RNPs) is a widespread strategy for regulating gene expression, yet how this process is orchestrated in mammalian systems remains unclear. Here, we show that Bicaudal D2 (BICD2), a partner of the microtubule motor dynein, is complexed with RNA-binding proteins (RBPs) in human cells. We also reveal that two of these RBPs – the Fragile X-related proteins FXR1 and FXR2 – bind directly to BICD2 and recruit mRNAs to dynein. Surprisingly, given its conventional designation as a cargo adaptor, BICD2 is not needed for dynein to interact with FXR1 and FXR2. Using *in vitro* reconstitution, we demonstrate that BICD2 instead activates dynein motility in concert with the FXR proteins. These data support a model in which BICD2 is recruited to pre-existing RNP-motor complexes, where it is licensed by the FXR proteins to activate processive movement. We further provide evidence that FXR proteins, BICD2 and dynein cooperate to promote rapid formation of stress granules, large RNPs that form in the cytoplasm in response to cellular insults. Collectively, our work reveals molecular mechanisms underlying RNP transport and stress granule assembly in human cells and indicates that cargo recruitment and motor activation are separable steps in dynein-mediated transport.

## INTRODUCTION

Microtubule motors play a central role in cellular organization through their ability to translocate organelles and macromolecules through the cytoplasm. Whereas a family of kinesin motors mediates cargo transport towards the plus ends of microtubules (Hirokawa *et al*, 2009), a single motor – cytoplasmic dynein-1 (hereafter dynein) – is tasked with moving cargoes towards the minus ends of these structures (Reck-Peterson *et al*, 2018). Dynein has evolutionarily conserved roles in transporting several membrane-bound compartments, including endosomes, lysosomes, mitochondria, and nuclei (Reck-Peterson *et al*, 2018). These functions depend on a set of coiled-coil-containing proteins, termed ‘activating adaptors’. Because these proteins bind simultaneously to dynein and proteins on the surface of specific cellular components, it is thought they play a key role in selectively recruiting cargoes to the motor complex (Olenick & Holzbaur, 2019; Reck-Peterson *et al*, 2018). In addition, activating adaptors switch on movement of the motor by stabilizing its interaction with the dynein-activating complex, dynactin (McKenney *et al*, 2014; Olenick & Holzbaur, 2019; Schlager *et al*, 2014). In this way, activating adaptors are proposed to directly link cargo engagement to long-range movement of motor complexes.

As exemplified by the female germline of the fruit fly *Drosophila melanogaster*, dynein can also translocate subsets of mRNAs through the cytoplasm to control where their protein products are synthesized and function. In the *Drosophila* system, mRNA transport is dependent on the Bicaudal D (BicD) activating adaptor and its binding partner Egalitarian (Egl), which engages structured elements within the RNA (Dienstbier *et al*, 2009; Singh *et al*, 2026). Docking of Egl relieves autoinhibition of BicD, triggering its association with dynein-dynactin and initiation of RNP transport (McClintock *et al*, 2018; Sladewski *et al*, 2018).

Dynein also mediates RNP trafficking in vertebrates, including in neurons (Ma et al, 2011; Tsai et al, 2009), oligodendrocytes (Herbert et al, 2017) and fibroblasts (Loschi et al, 2009). However, as Egl is restricted to invertebrates, the factors that link RNPs to dynein in vertebrate cells remain unclear. In contrast, BicD proteins are present throughout the animal kingdom, with mammals possessing two paralogs. These proteins – BICD1 and BICD2 – have well-established roles in dynein-based transport of Golgi-derived vesicles, nuclei, centrosomes and pathogens (Hoogenraad & Akhmanova, 2016; Manigrasso *et al*, 2025).

In this study, we set out to define proteins that participate in dynein-based RNP transport in human cells. We show that BICD2 is complexed with RBPs, including FXR1 and FXR2, to which it binds directly. We also demonstrate that the FXR proteins link mRNAs to dynein. Despite its interactions with FXR1 and FXR2, BICD2 does not serve as an obligatory linker between these proteins and the motor complex. Our data indicate that BICD2 instead functions with the FXR proteins to activate dynein motility. Further, we provide evidence that dynein, BICD2 and the FXR proteins cooperate to promote assembly of stress granules, large cytoplasmic RNPs that form in response to cellular stressors and modulate translation of associated mRNAs (Protter & Parker, 2016; Smith & Bartel, 2026). Collectively, our study identifies machinery for RNP transport and stress granule assembly in mammalian cells and challenges the widely accepted view of activating adaptor function in dynein-based cargo transport.

## RESULTS

### BICD2 and dynein are complexed with RBPs in human cells

We reasoned that, although Egl is not present in mammals, the role of BicD proteins as activating adaptors during dynein-based mRNA transport could be conserved. To explore this possibility, we searched for BICD1 and BICD2 in the R-DeeP database, which identifies proteins whose sedimentation profiles in HeLa cell extracts are shifted by RNase-treatment, indicative of incorporation into RNA-containing complexes (Caudron-Herger *et al*, 2019). BICD1 was not represented in the database either in the presence or absence of RNase, potentially due to its relatively low abundance in HeLa cells (Nagaraj *et al*, 2011). However, a substantial fraction of BICD2 was found in species that decreased in apparent size upon RNase treatment (Figure S1), consistent with its presence in RNPs. None of the other known activating adaptors for dynein exhibited RNase-sensitive sedimentation profiles in the R-DeeP database.

To shed further light on the composition of complexes containing BICD2, we used a GFP-binding nanobody to precipitate GFP-tagged BICD2 from HeLa cells that were grown under basal conditions. The captured material was then analyzed by label-free quantitative mass-spectrometry to identify co-purifying proteins. Proteins captured with GFP-BICD2 were compared to those isolated using the same procedure from cells expressing either a GFP-tagged version of the dynein subunit DYNC1I2 or GFP alone. This approach was designed to reveal RBPs that selectively associate with both BICD2 and dynein, and are therefore candidates to participate in mRNA transport. In parallel, to assess the RNA dependence of interactions with BICD2 and dynein, we performed equivalent experiments using extracts in which RNA was degraded with RNase.

Supporting the validity of our approach, the proteins enriched with GFP-BICD2 versus GFP in the absence of RNase included dynein components and another known partner of BICD2, the kinase PLK1 (Gallisa-Sune *et al*, 2023) (Figure 1A). Strikingly, many of the other BICD2 interactors in this condition were known RNA-associated proteins (Figure 1A–C). The majority of these proteins were also selectively captured with GFP-DYNC1I2 in the RNase-free samples (Figure 1B, Figure S2A, B). For example, both BICD2 and DYNC1I2 associated with components of the 60S large ribosomal subunit (Figure 1A, B). Constituents of the 40S small ribosomal subunit were not recovered with either protein, which we confirmed in separate experiments (Figure S2C) was due to splitting of the ribosomal subunits by the presence of ethylenediaminetetraacetic acid (EDTA) in our assay buffer (Sabatini *et al*, 1966). These data reveal that both BICD2 and DYNC1I2 complex with intact ribosomes via the large ribosomal subunit.

**Figure 1:**
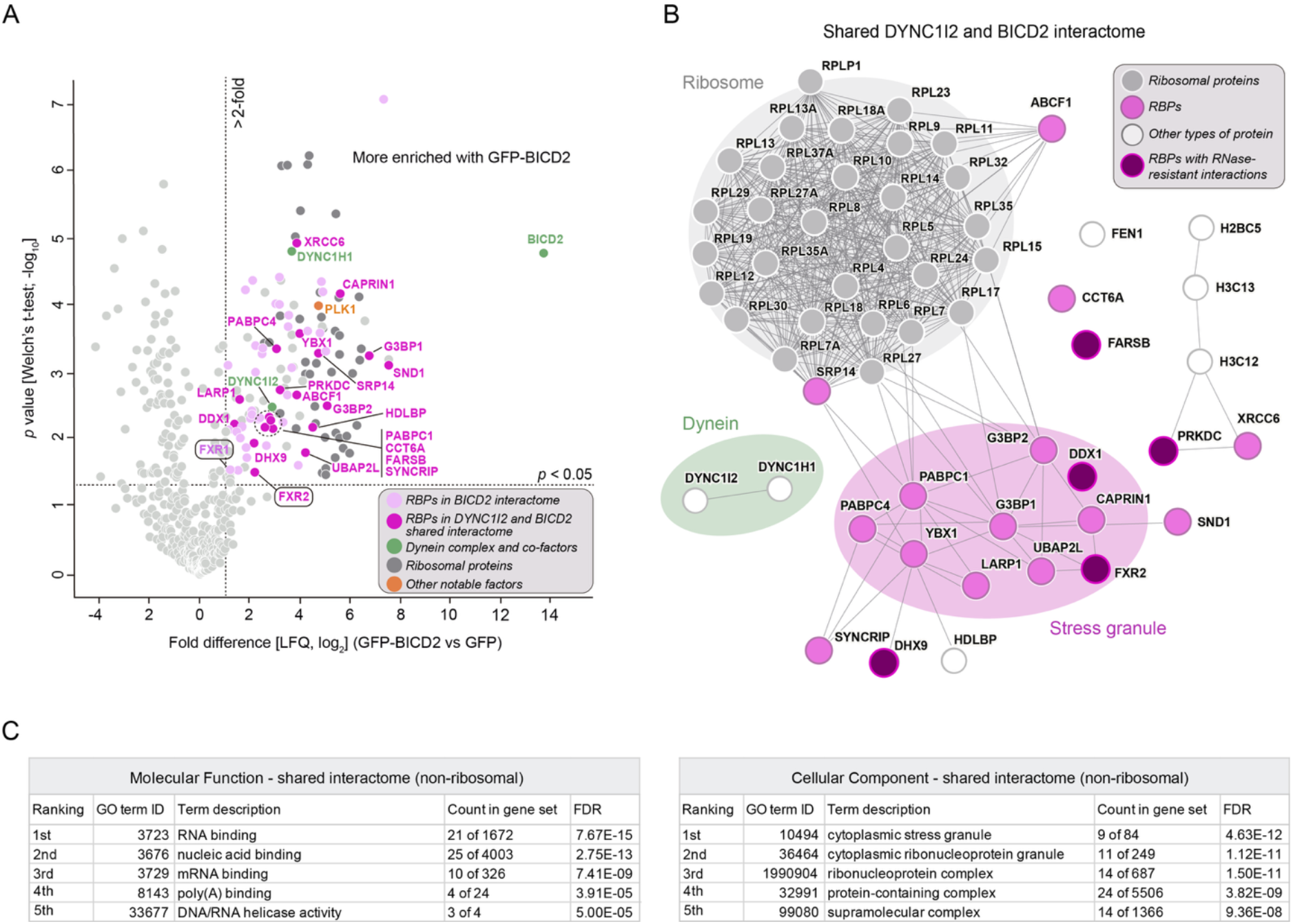
BICD2 and dynein associate with a shared set of RBPs. **(A)** Volcano plot showing log_2_-fold enrichment of proteins in GFP-BICD2 versus GFP immunoprecipitates from HeLa cells, as well as associated *p* values (three biological replicates per condition). LFQ, label-free quantification. **(B)** STRING analysis of proteins significantly enriched (> 2-fold enrichment and *p* < 0.05) in both GFP-DYNC1I2 and GFP-BICD2 immunoprecipitates relative to their respective GFP only controls. Connecting lines indicate previous experimental evidence of complexation between these proteins. RNase resistance was defined as a log_2_ fold change in association upon RNase treatment of < −1.58 (corresponding to a reduction of <66.7%) in both interactomes. **(C)** Gene ontology (GO) analysis of the shared DYNC1I2 and BICD2 interactome, excluding ribosomal proteins. As FXR2 was not classified as a stress granule protein in the GO database, despite experimental evidence that this is the case (Curdy *et al*, 2023), only nine stress granule components were highlighted by this analysis. FDR, false discovery rate.

Both BICD2 and DYNC1I2 additionally associated with 19 non-ribosomal RBPs (Figure 1B). Notably, many of these factors – CAPRIN1, DDX1, FXR2, G3BP1, G3BP2, LARP1, PABPC1, PABPC4, UBAP2L and YBX1 – are known constituents of stress granules. Together, these data show that dynein and BICD2 share many interactors that are components of RNA-containing complexes. Thus, BICD2 is a strong candidate to participate in dynein-based RNP transport.

Intriguingly, BICD2 was not detected in the DYNC1I2 interactome (Figure 1B and Figure S2A). The finding that dynein can be complexed with the same RBPs as BICD2 even when the latter protein is not detected raises the possibility that BICD2 is not a constitutive component of RNP-dynein complexes. Such a scenario would be incompatible with BICD2 acting as an obligatory linker between RBPs and dynein and would instead be consistent with a regulatory role for this protein in RNP trafficking.

We next examined the effect of RNase on the interactomes of BICD2 and DYNC1I2. RNase treatment substantially impaired the association of ribosomal proteins with BICD2, but not with DYNC1I2 (Figure S3A, B). The observation that dynein remains complexed with ribosomes under conditions where BICD2 is depleted further supports the notion that BICD2 is not essential for linkage of the motor to RNPs.

The interactions of both BICD2 and DYNC1I2 with most other RNA-associated proteins were strongly reduced by RNase (Figure 1B and Figure S3A, B). In contrast, BICD2 and DYNC1I2 exhibited a relatively modest reduction in association with FXR2 following RNase treatment, while their co-purification with the RNA helicases DDX1 and DHX9, the tRNA ligase FARSB, and the nucleic-acid binding kinase PRKDC was either unaffected or enhanced by RNA degradation (Figure 1B and Figure S3A, B). Collectively, our proteomic analyses reveal RBPs that associate with both BICD2 and dynein, and distinguish interactions that are strongly RNA-dependent from those that are resistant to RNA degradation.

### FXR proteins are candidate RNA adaptors for BICD2 and dynein

Proteins whose associations with BICD2 and dynein are strongly impaired by RNase treatment may associate indirectly with the transport machinery by ‘hitchhiking’ on the RNA cargo. We therefore focused on the RBPs whose interactions were either unimpaired, or only modestly affected, by RNase as candidate RNA-binding adaptors for the motor complex. To narrow down these candidates further, we asked if they were recovered in a BioID proximity-labeling screen using the cargo-binding third coiledcoil domain (CC3) of BICD2 in human HEK293 cells (Redwine *et al*, 2017). Only FXR2 met this criterion and was therefore prioritized for further study. We also selected FXR1, a closely related paralog of FXR2 that also localizes to stress granules (Garvanska *et al*, 2024; Sanders *et al*, 2020; Say *et al*, 2010), for additional investigation because it was present in our BICD2 interactome (Figure 1A) and narrowly missed the threshold for classification as a DYNC1I2 interactor (Figure S2A).

Together with the fragile X mental retardation protein FMRP, which was absent in both the BICD2 and DYNC1I2 interactomes, FXR1 and FXR2 constitute the FXP family. Each of these proteins contains RNA-binding K-homology (KH) and RGG domains, and has been implicated in multiple post-transcriptional events, including splicing, RNA editing and translational control (Majumder et al, 2020). Analysis of additional nanobody capture experiments by immunoblotting demonstrated that both FXR1 and FXR2 co-purify with BICD2 and DYNC1I2 from HeLa cells, including when RNA is degraded (Figure 2A, B). We also found that BICD2, as well as the dynactin component DCTN1, were selectively enriched in the material immunoprecipitated with GFP-FXR2 from HeLa cells (Figure 2C). Supporting the results of the immunoprecipitation experiments, *in situ* proximity ligation assays (Soderberg *et al*, 2006) revealed that both FXR1 and FXR2 are closely apposed to BICD2 in the cytoplasm of HeLa cells (Figure 2D). Together, these data confirm that FXR proteins can complex with BICD2 and the dynein transport machinery.

**Figure 2:**
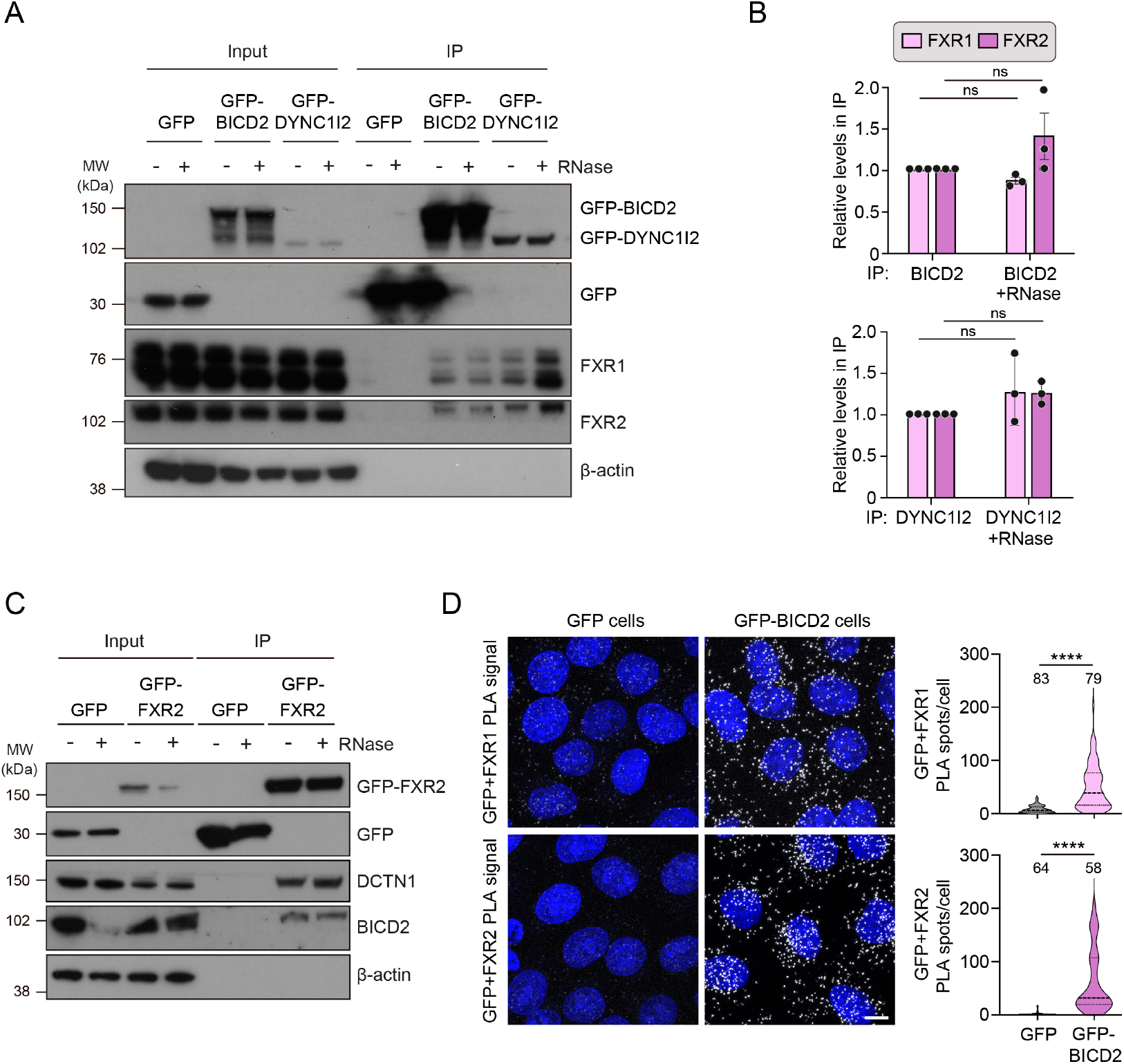
Validation of association of FXR proteins with BICD2. **(A)** Immunoblots assessing presence of indicated proteins in GFP immunoprecipitates from GFP-BICD2, GFP-DYNC1I2 or GFP HeLa cell lysates treated with RNase (+) or RNase inhibitor (-). MW: molecular weight markers. IP: immunoprecipitated material. In A and C, β-actin serves as a negative control, demonstrating specificity of the immunoprecipitation procedure. **(B)** Quantification of FXR1 or FXR2 signals in independent immunoprecipitation-immunoblot experiments (N = 3). Circles represent per experiment values normalized to both the bait and the corresponding RNase inhibitor condition; error bars show standard deviation (SD). **(C)** Immunoblots assessing presence of indicated proteins in GFP immunoprecipitates from GFP-FXR2 or GFP HeLa cell lysates treated with RNase (+) or RNase inhibitor (-). **(D)** Confocal images (left) and quantification (right) from proximity ligation assay (PLA) experiments in HeLa cells expressing GFP only or GFP-BICD2 and using primary antibodies to GFP and either FXR1 or FXR2. White spots in the images indicate molecular proximity of the protein targets. Scale bar: 10 μm. In the violin plots, solid lines and dashed lines denote medians and interquartile range, respectively; numbers of cells analyzed are shown above the plots. In B and D, statistical significance was evaluated with a two-tailed unpaired t-test (B) or a Krustal-Wallis test (D) (ns, not significant; ****, *p* < 0.0001).

### FXR proteins bind directly to the cargo-binding region of BICD2

The finding that BICD2 is found in a complex with FXR1 and FXR2 even when RNA is degraded raised the possibility that BICD2 can interact directly with these proteins. We showed that this is indeed the case by coupling recombinant BICD2 to beads and performing pulldown assays with recombinant FXR1 or FXR2 proteins (Figure 3A). To shed light on the determinants of BICD2-FXR interactions, we performed additional pulldown assays with FXR1 and truncated versions BICD2. This revealed that the C-terminal region of BICD2 that contains the CC3 domain (BICD2^CC3^) is sufficient to bind FXR1 (Figure 3B), whereas truncations lacking an intact CC3 do not interact with this protein (Figure S4A, B). This observation is consistent with the well-established role of CC3 in binding cargo-associated proteins (Hoogenraad & Akhmanova, 2016). Corroborating these *in vitro* observations, immunoprecipitations of GFP-tagged proteins from HeLa cells showed that the BICD2^CC3^ region is sufficient to associate with FXR1 and FXR2 *in vivo* (Figure S4C).

**Figure 3:**
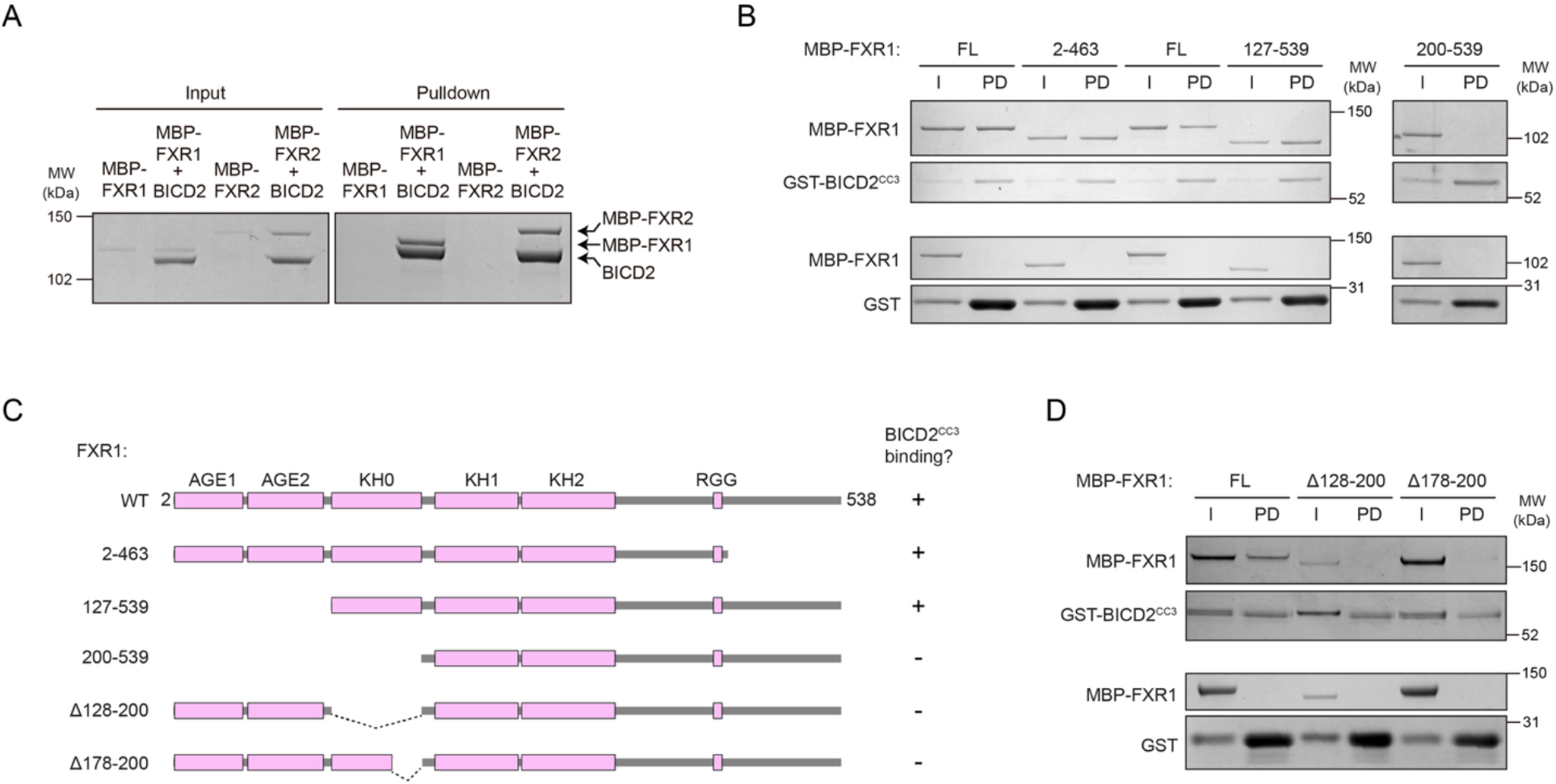
BICD2 interacts directly with FXR proteins. **(A)** Images of Coomassie-stained gels showing association of recombinant MBP-tagged human FXR1 or FXR2 with mouse BICD2 in pulldown assays; strep-tagged BICD2 was used as bait. The C-terminal unstructured region of each FXR protein was removed to enable production of soluble protein (Edwards *et al*, 2020). MW: molecular weight markers. **(B)** Images of Coomassie-stained gels revealing interactions of truncated forms of FXR1 with BICD2^CC3^ (residues 577–820 of the mouse sequence). I, input; PD, pulldown. **(C)** Summary of data from pulldown assays with GST-BICD^CC3^ and MBP-FXR1 variants. Folded domains are shown in magenta. WT, wild type. **(D)** Images of Coomassie-stained gels assaying interactions of BICD2^CC3^ with FXR1 variants that have the indicated KH0 domain deletions. In B–D, GST-BICD^CC3^ (or GST only for the specificity control) was used as bait.

We next asked if the region of BICD2^CC3^ (amino acids (aa) 711–800) that binds known cargo-associated partners – including the Golgi vesicle adaptor RAB6, the nucleoporin RANBP2/NUP358, and the bacterial surface protein ScaC (Gibson *et al*, 2023; Hoogenraad & Akhmanova, 2016; Manigrasso *et al*, 2025) – is sufficient to engage FXR proteins. This protein did not bind FXR1 *in vitro* (Figure S4A, B), suggesting a non-canonical mode of interaction with BICD2. Consistent with this finding, mutation of a CC3 residue (L786 mouse/L790 human) that is required for interaction of BICD2 with the prototypical cargo-associated partners RAB6 and Egl (Liu *et al*, 2013; Manigrasso *et al*, 2025) did not impair engagement with FXR1 or FXR2 in HeLa cells (Figure S4C).

To elucidate which region of FXR proteins is responsible for BICD2 binding, we performed *in vitro* pulldown assays using truncated versions of FXR1. Comparing the results obtained with FXR1^127–539^ and FXR1^200–539^ (Figure 3B, C) revealed that residues 127–199 of FXR1 are essential for association with BICD2^CC3^. This region of FXR1 corresponds to the KH0 domain, which, unlike the other KH domains in FXP family members, lacks a canonical RNA-binding motif (Myrick *et al*, 2015). Structural modeling with AlphaFold 3 (Abramson *et al*, 2024) consistently predicted an interaction between the C-terminal portion of the KH0 domains of FXR1 (aa 176–201) and FXR2 (aa 189–211) with positions 734–761 of BICD2^CC3^ (Figure S5A, B). Whilst the inability of BICD2^711–800^ to bind FXR1 (Figure S4A) indicates that the predicted interface is not sufficient for a stable BICD2-FXR interaction, its importance for binding is supported by our observation that removing residues 128–200 or 178–200 of FXR1 strongly reduced association with BICD2^CC3^ (Figure 3C, D). This result is unlikely to reflect misfolding of FXR1 when the KH0 domain is perturbed, as differential scanning fluorimetry revealed indistinguishable thermal denaturation profiles of wild-type and Δ178-200 FXR1 (Figure S5C). Taken together, these data reveal direct binding of BICD2 to FXR proteins and shed light on molecular determinants of this interaction.

### FXR1 and FXR2 connect mRNAs to BICD2 and dynein

The finding that FXR1 and FXR2 engage the cargo-binding region of BICD2 is consistent with them acting as adaptors between mRNAs and BICD2-containing dynein transport complexes. If this were the case, one would expect depletion of the FXR proteins to reduce the association of mRNAs with BICD2 and dynein. To investigate this possibility, we immunoprecipitated GFP-BICD2 or GFP-DYNC1I2 from HeLa cells that had been pre-treated with either an siRNA pool that depletes both FXR proteins (siFXR1+2; Figure S6A) or a non-targeting siRNA control pool (siCTRL). RT-qPCR was then used to compare the abundance of recently identified FXR1 target mRNAs (Chen *et al*, 2024) in the captured material.

In the presence of the siRNA control, all three FXR1 target mRNAs analyzed – *G3BP1, HSPA8*, and *TUBB* – were significantly enriched in the GFP-BICD2 and GFP-DYNC1I2 samples relative to samples from cells expressing GFP only (Figure 4). We also found that the degree of enrichment of each of these mRNAs with GFP-BICD2 or GFP-DYNC1I2 over the GFP control was significantly reduced by FXR1 and FXR2 depletion (Figure 4). This effect was not attributable to the FXR knockdown lowering the overall abundance of target mRNAs in the cell, as bulk RNA sequencing of extracts showed that *FXR1* and *FXR2* were the only transcripts that were substantially depleted by the FXR siRNAs (Figure S6B, C). These observations demonstrate that FXR1 and FXR2 promote the association of mRNAs with BICD2 and dynein and are therefore *bona fide* RNA adaptors for the transport machinery.

**Figure 4:**
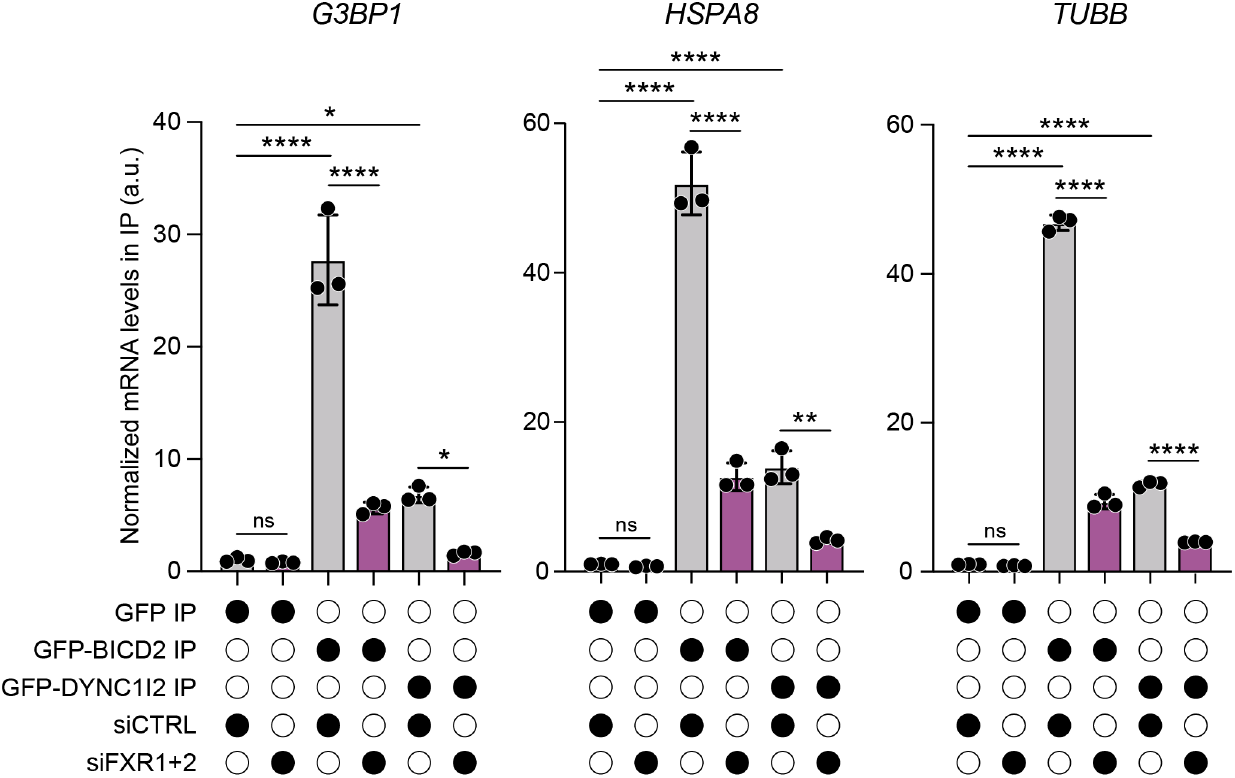
FXR proteins couple mRNAs to BICD2 and dynein. Abundance of indicated FXR1 target mRNAs in GFP immuno-precipitates from GFP, GFP-BICD2 or GFP-DYNC1I2 HeLa cells treated with the indicated siRNA pools, as determined by RT-qPCR. Values are normalized to the siCTRL values from GFP only cells. Small black circles are values for individual experiments. Large black circles indicate experimental conditions. Statistical significance was evaluated with a one-way ANOVA test with Tukey’s multiple comparisons correction (N = 3; ns, not significant; *, *p* < 0.05; ****, *p* < 0.0001).

### FXR proteins and BICD2 have distinct roles in dynein recruitment and activation

As described above, the canonical model for activating adaptor function assigns these proteins two roles: linking cargo to dynein and activating motor motility (Olenick & Holzbaur, 2019; Reck-Peterson *et al*, 2018). However, our detection of FXR2, but not BICD2, in DYNC1I2 immunoprecipitates (Figure S2A) raised the possibility that BICD2 is not obligatory for linkage of mRNA adaptors to dynein. To test this notion further, we used immunoprecipitation to determine how the association of FXR1 and FXR2 with GFP-DYNC1I2 is affected by an siRNA pool that strongly depletes BICD2 (siBICD2) (Figure S6A). Knockdown of BICD2 did not disrupt the interaction of FXR1 and FXR2 with dynein (Figure 5A). This was not due to functional redundancy between BICD1 and BICD2, as simultaneous depletion of both BICD proteins (siBICD1+2) (Figure S6A) also did not impair interaction of the FXR proteins with the motor complex (Figure 5A). These observations demonstrate that BICD2 is not required for linkage between the FXR proteins and dynein. Consistent with this notion, we detected direct interactions between the recombinant human dynein complex and recombinant FXR1 and FXR2 in an *in vitro* pulldown assay (Figure S7).

**Figure 5:**
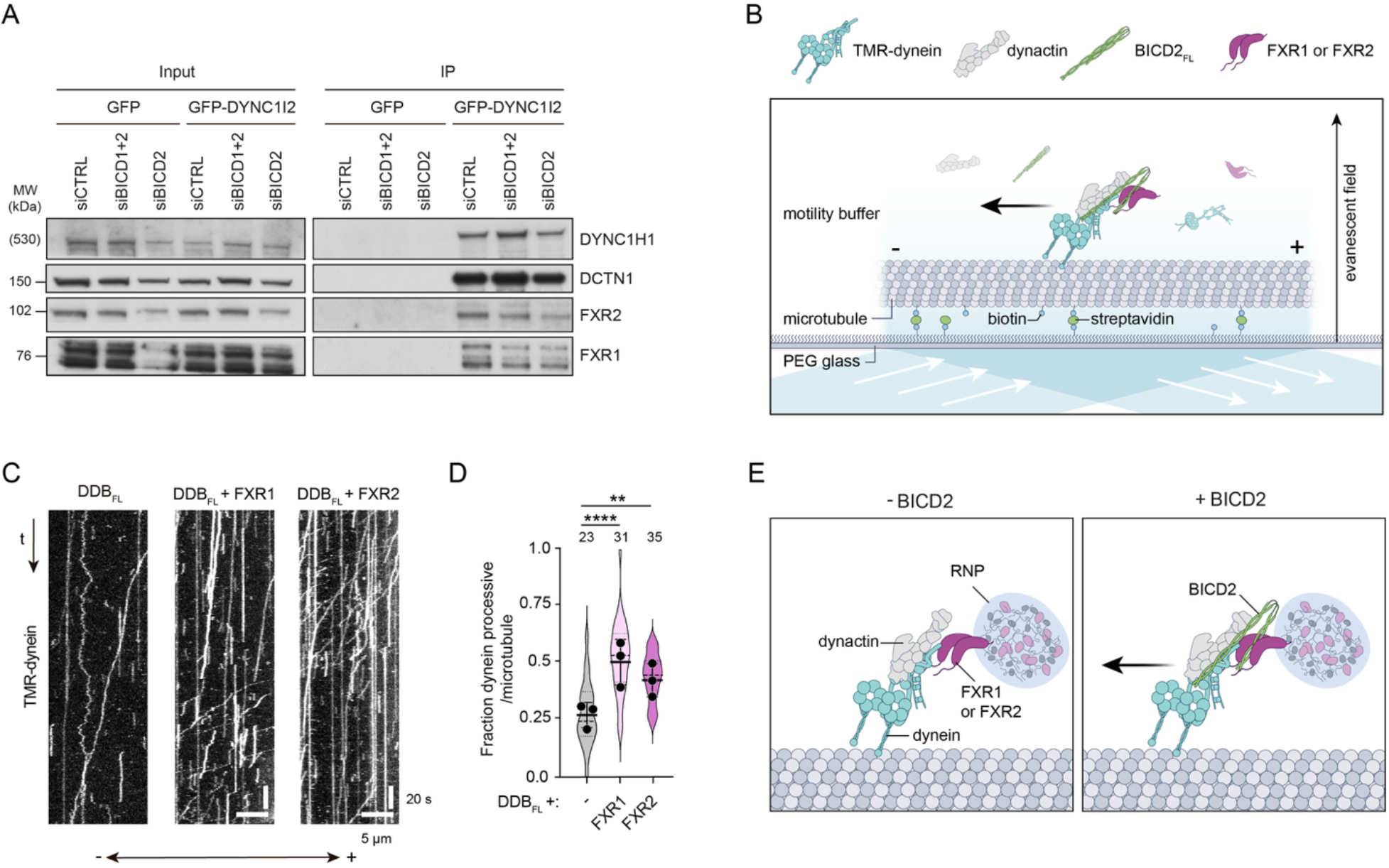
FXR proteins associate with dynein independently of BICD2 and enhance BICD2-mediated motor activation. **(A)** Immunoblots assessing the presence of indicated proteins in GFP immunoprecipitates from GFP or GFP-DYNC1I2 HeLa cells treated with the indicated siRNA pools. (B) Cartoon of *in vitro* motility assay, which uses combinations of full-length (FL) BICD2, FXR1, FXR2, dynactin and Tetramethylrhodamine (TMR)-labeled dynein. MT, microtubule; PEG, polyethylene glycol. (C) Example kymographs (time-distance plots) of TMR-dynein behavior in the presence of the indicated factors. DDB_FL_, dynein-dynactin-BICD_FL_; t, time. (D) Quantification of fraction of microtubule-bound dyneins that exhibit processive movement. Violin plots represent data for individual microtubules, with circles representing mean values from individual experiments (N = 3) and error bars representing SD. Solid and dashed lines show median values and interquartile range, respectively. Numbers of microtubules are shown above bars. Statistical significance was evaluated with a one-way ANOVA test with Tukey’s post-hoc multiple comparisons correction using per microtubule values aggregated across experiments (**, *p* < 0.01; ****, *p* < 0.0001). (E) Model for role of BICD2 in FXR-based RNP transport. Arrow signifies stimulation of motility. Complexation of dynactin with FXR proteins and dynein when BICD2 is absent is hypothetical. However, this scenario is consistent with recent structures showing dynein-dynactin association on microtubules in the absence of activating adaptors (Rao *et al*, 2026) and our observation that dynactin subunits and FXR2, but not BICD2, are detected in DYNC1I2 immunoprecipitates (Figure S2A).

These observations led us to hypothesize that BICD2 acts primarily to stimulate motility of RNPs that are bound by FXR proteins and dynein. In other cargo transport pathways, binding of cargo-associated proteins to CC3 relieves autoinhibition of BicD family members, allowing their N-terminal regions to engage and activate dynein-dynactin (Hoogenraad *et al*, 2003; Liu *et al*, 2013; Terawaki *et al*, 2015). Because FXR proteins interact with BICD2^CC3^ (Figure 3B), we reasoned that they might promote dynein-based transport in an analogous manner.

To test if BICD2 can stimulate dynein motility in conjunction with FXR proteins, we attempted to reconstitute this process *in vitro* using purified proteins. Full-length BICD2 (BICD2_FL_) was mixed with dynactin and fluorescently-labeled dynein in the presence and absence of FXR1 or FXR2. These mixtures were then diluted and introduced into a chamber that had fluorescent microtubules immobilized on a glass surface (Figure 5B), enabling visualization of transport-competent complexes by total internal reflection fluorescence (TIRF) microscopy.

As expected from the autoinhibited nature of cargo-free BICD2 (Hoogenraad & Akhmanova, 2016; Huynh & Vale, 2017), there was limited transport of dynein when it was incubated with only dynactin and BICD2_FL_ (Figure 5C, D). In contrast, combining either of the FXR proteins with BICD2_FL_ and dynactin significantly increased the proportion of microtubule-bound dynein that underwent active transport (Figure 5C, D). The FXR proteins also increased the incidence of microtubule binding by dynein in the presence of BICD2_FL_ and dynactin (Figure S8A), reminiscent of the enhanced microtubule engagement conferred by dynactin and a truncated version of BICD2 that is constitutively active due to a lack of autoinhibition (Belyy *et al*, 2016). FXR2, but not FXR1, also caused a modest increase in dynein velocity in the presence of BICD2 and dynactin without significantly affecting run length (Figure S8A).

Importantly, when BICD2_FL_ was omitted, neither FXR1 nor FXR2 was sufficient to activate dynein movement or microtubule binding in the presence of dynactin (Figure S8B). This is consistent with the critical role of activating adaptors in licensing motility of the motor complex in previous studies (McKenney *et al*, 2014; Olenick & Holzbaur, 2019; Schlager *et al*, 2014).

Collectively, these results show that FXR proteins work together with BICD2 to stimulate dynein-based transport. Our findings support a model in which BICD2 is recruited to complexes containing FXR proteins and dynein, with FXR binding relieving BICD2 autoinhibition to switch on motor movement (Figure 5E).

### FXR and BICD proteins promote stress granule assembly

Finally, we turned our attention to the functional significance of the FXR-BICD2-dynein interaction. Despite being generated using cells cultured in basal conditions, our shared BICD2 and DYNC1I2 interactome contained several stress granule components (Figure 1B). We therefore hypothesized that FXR proteins, BICD2 and dynein cooperate to promote the formation of stress granules when cells experience an insult, potentially by bringing together RNP ‘seeds’ via transport along microtubules. In line with this possibility, two of the three mRNAs for which we demonstrated an FXR-dependent interaction with BICD2 and dynein (*G3BP1* and *HSPA8*; Figure 4) have been reported to be enriched in stress granules (Khong *et al*, 2017; Ren *et al*, 2023).

Consistent with a role for FXR proteins and BICD2 in promoting assembly of stress granules, immunostaining of U2OS cells revealed association of all three proteins with these structures following their induction by sodium arsenite, a commonly used oxidative stressor (Figure 6A and Figure S9A).

**Figure 6:**
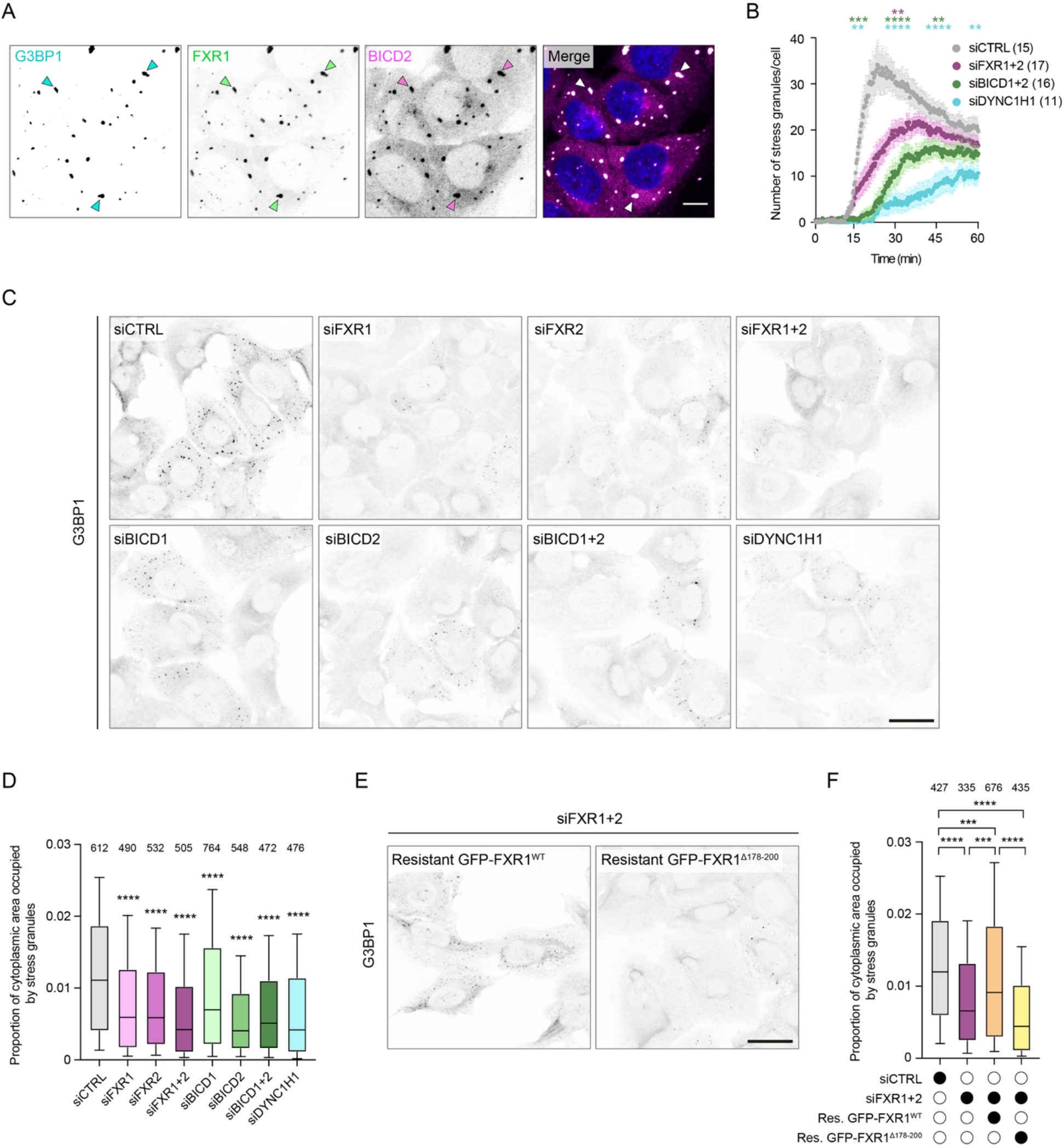
FXR and BICD proteins promote stress granule assembly. **(A)** Confocal images of immunostained U2OS cells that have been treated with 300 μM arsenite for 1 h. Arrowheads show examples of stress granules. **(B)** Time course of stress granule formation (monitored by GFP-G3BP1) in U2OS cells treated with the indicated siRNA pools for the previous 72 h and exposed to 300 μM arsenite at 0 min. Circles and error bars denote mean and SD, respectively. Numbers of cells per condition are shown in parentheses. **(C)** Confocal images of stress granules (marked with an antibody to endogenous G3BP1) in U2OS cells treated with the indicated siRNA pools and fixed 20 min after arsenite addition. **(D)** Quantification of stress granule formation in the experiment documented in C. **(E)** Confocal images of stress granules (marked with G3BP1 antibodies) in cells expressing genomically-integrated siRNA-resistant GFP-tagged wild-type (WT) or Δ178–200 FXR1 transgenes and pretreated with siRNAs targeting endogenous FXR1 and FXR2. Cells were exposed to 300 μM arsenite for 20 min. **(F)** Quantification of stress granule formation in the indicated conditions. Res., siRNA-resistant. In D and F, horizontal lines denote median, boxes denote interquartile range and whiskers denote 10^th^-90^th^ percentile values. Number of cells per condition (from N = 4 (D) or 3 (F) independent experiments) is given above bars. In B, D and F, statistical significance compared to siCTRL was evaluated with a one-way ANOVA test with Tukey’s multiple comparisons correction. In B, datasets were compared at 15, 30, 45 and 60 min, with asterisks colored according to which condition is compared to siCTRL. *, *p* < 0.05; **, *p* < 0.01; ***, *p* < 0.001; ****, *p* < 0.0001. Scale bars represent 10 μm (A) and 30 μm (C and E).

We next assessed the functional contribution of FXR proteins, BICD2 and dynein to arsenite-induced assembly of stress granules. This was done by siRNA-mediated knockdown of these factors in U2OS cells that expressed a GFP-tagged version of the stress granule marker G3BP1. Live imaging revealed significant delays in formation of stress granules following treatment with siRNAs targeting Dynein heavy chain (siDYNC1H1), FXR1 and FXR2 (siFXR1+2), or BICD1 and BICD2 (siBICD1+2) (Figure 6B and Movie S1). Impaired stress granule formation in the knockdown cells was most evident within the first 30 minutes of arsenite addition (Figure 6B), indicating that dynein, FXR and BICD proteins are particularly important during the early stages of stress granule assembly.

As these experiments used siRNA pools that target both FXR paralogs or both BICD2 paralogs simultaneously, we next determined the effects of depleting each protein individually. By immunostaining U2OS cells that were fixed 20 minutes after arsenite addition, we found that separate knockdowns of FXR1, FXR2, BICD1 and BICD2 significantly impaired stress granule formation, although targeting BICD1 had the weakest effect (Figure 6C, D and Figure S9B). We additionally used this assay to corroborate our previous observation that siRNA-mediated depletion of DYNC1H1 compromises assembly of stress granules (Figure 6C, D).

To explore whether FXR proteins and BICD2 could function together to promote stress granule assembly, we assessed the consequences of disrupting their biochemical interaction. We found that an siRNA-resistant wild-type FXR1 transgene suppressed the stress granule assembly phenotype caused by simultaneous knockdown of endogenous FXR1 and FXR2 (Figure 6E, F and Figure S9C). In contrast, a version of the FXR1 transgene that lacks the region required for BICD2 binding (FXR1^Δ178-200^; Figure 3D) was unable to alleviate this defect (Figure 6E, F and Figure S9C), despite being expressed at comparable levels to its wild-type counterpart (Figure S9D). Together, our findings indicate that FXR proteins and BICD2 cooperate with each other and dynein to promote assembly of stress granules.

## DISCUSSION

As the major minus-end-directed microtubule motor in most eukaryotic cells, dynein is responsible for translocating a wide variety of cellular constituents. The mechanisms by which dynein traffics its diverse cargoes are only partly understood. However, a class of motor co-factors known as activating adaptors are known to play a central role in this process. These proteins stimulate dynein’s motility by stabilizing its association with the processivity factor dynactin (McKenney *et al*, 2014; Olenick & Holzbaur, 2019; Schlager *et al*, 2014) and have additionally been proposed to recruit motors to specific cargoes through interactions with cargo-associated proteins (Olenick & Holzbaur, 2019; Reck-Peterson *et al*, 2018). According to this framework, activating adaptors attach cargoes to dynein and concomitantly trigger motility of the motor complex. However, the extent to which this model applies across the diverse range of dynein cargoes is not clear. Moreover, for several cargo classes, including mammalian RNPs, the factors that mediate cargo recruitment and motor activation have not been identified.

Here, we show that BICD2, an activating adaptor that was previously shown to promote dynein-based transport of Golgi vesicles, nuclei and pathogens, associates with RNPs in human cells. We also reveal that BICD2 binds directly to FXR1 and FXR2, and that these RBPs recruit mRNAs to dynein. Contrary to prevailing models of activating adaptor function, we show that BICD2 is dispensable for linkage of FXR proteins to dynein and serves primarily as an activator of motor movement. Our *in vitro* data indicate that the FXR proteins can interact directly with the dynein complex, offering an explanation for why BICD2 is not required for a connection between these factors. Taken together, our data support a model in which BICD2 is recruited to pre-existing FXR-dynein complexes, with FXR engagement relieving BICD2 autoinhibition and thereby activating dynein motility.

Uncoupling cargo linkage and motor activation might be advantageous as it allows these events to be regulated independently, thereby conferring greater spatial and temporal control over trafficking. Having dynein recruitment precede motor activation may be particularly beneficial during transport of RNPs, as these structures often need to mature via compositional remodeling (Tauber *et al*, 2020). Recruitment of dynein to FXR-bound RNPs would create a primed but inactive transport complex, with later engagement of BICD2 with FXR proteins and dynein acting as a switch that initiates processive movement once a mature RNP is formed. The regulation of pre-recruited motors by activating adaptors could extend beyond RNP trafficking. This possibility is supported by the lack of correlation between levels of BicD and dynein on lipid droplets during *Drosophila* development (Larsen *et al*, 2008), as well as reports of direct interactions between organelle-bound partners of activating adaptors and dynein-dynactin components (Johansson *et al*, 2007; Short *et al*, 2002).

Whereas FXR proteins have previously been shown to cooperate with BICD2 to control the formation of nucleoporin condensates (Agote-Aran *et al*, 2020), to our knowledge they have not previously been implicated in RNP transport. In *Drosophila*, the sole member of the FXP family, FMRP, was previously found in a complex with BicD (Bianco *et al*, 2010), the fly ortholog of BICD1 and BICD2. However, the molecular function of this interaction was not resolved. Intriguingly, the nature of the interaction between FXP and BicD proteins appears to differ in flies and mammals. Binding of *Drosophila* FMRP to BicD depends on the integrity of the canonical cargo-binding site on BicD^CC3^ that is used by RAB6 and Egl, and additionally requires the presence of RNA (Bianco *et al*, 2010). In contrast, BICD2 engages FXR1 and FXR2 independently of the canonical docking site, including when RNA is absent. These observations suggest that interactions of FXP family members with BicD proteins have been functionally adapted in flies and mammals.

Structural modeling offers further insight into the molecular basis of the distinct FXR-BICD2 interaction in mammalian cells. AlphaFold 3 predicts that FXR1 and FXR2 bind to a region of BICD2^CC3^ that partially overlaps with a non-canonical interaction site for RANBP2/NUP358 (Gibson *et al*, 2023; Manigrasso *et al*, 2025). This raises the possibility that FXR proteins couple mRNAs to BICD2 without competing with cargoes that bind the canonical site. This arrangement could conceivably allow co-transport of RNPs and at least some membrane-bound cargoes, a phenomenon observed in several polarized cell types (Vargas *et al*, 2022). The presence of multiple cargo-binding sites on BICD2 might also permit simultaneous docking of different RBPs, which could increase the specificity of mRNA selection through combinatorial recognition of *cis*-acting elements. Intriguingly, our pulldown experiments indicate that the predicted interface between FXR and BICD2’s CC3 domain is necessary but not sufficient for stable association of these proteins. Defining additional interaction sites between these proteins and determining how FXR1 and FXR2 bind dynein independently of BICD2 will be priorities for future work.

In addition to shedding light on the interaction between FXR proteins and BICD2 and their division of labor during cargo recruitment and dynein activation, our findings point to a cellular function of the FXR-BICD2-dynein complex: promoting the rapid assembly of stress granules. Although recent work on stress granule formation has focused on the role of liquid-liquid phase separation (LLPS) (e.g., Guillen-Boixet *et al*, 2020; Lieber *et al*, 2026; Lindstrom *et al*, 2022; Molliex *et al*, 2015; Yang *et al*, 2020), two early studies provided evidence for an involvement of dynein (Loschi *et al*, 2009; Tsai *et al*, 2009). Loschi *et al* additionally reported that BICD1, but not BICD2, promotes stress granule assembly. In contrast, we find that both BICD1 and BICD2 contribute to this process. These contrasting observations may reflect differential deployment of BicD family members in different cell types or species, as Loschi and colleagues used mouse and monkey fibroblasts, whereas we studied human cancer cells. Importantly, our findings further extend the earlier observations of dynein’s involvement in stress granule assembly by linking the FXR-BICD2 interaction to this process.

We wish to emphasize that our data do not rule out a critical role for LLPS in the biogenesis of stress granules. In fact, we favor a model in which phase separation and dynein-based transport make complementary contributions to this process. This view is supported by our observation that depletion of FXR or BICD proteins substantially delays, but does not abolish, stress granule formation. This is consistent with stress granules being detectable in U2OS cells in which *FXR1* and *FXR2* (as well as *FMRP*) have been simultaneously knocked out (Sanders *et al*, 2020). We hypothesize that dynein-based motility facilitates encounters between dispersed RNP ‘seeds’, whose diffusion in the cytoplasm is expected to be strongly constrained by molecular crowding in the cytoplasm (Luby-Phelps *et al*, 1986). By promoting such encounters, dynein-driven translocation along microtubules would facilitate the formation of nascent clusters of these stress granule precursors. LLPS could then amplify these initial assemblies through further recruitment and concentration of RNAs and proteins, resulting in the production of mature stress granules.

Dynein-mediated nucleation of stress granules could conceivably be triggered by stress-induced signaling pathways that increase the frequency or range of movements mediated by FXR-BICD2 complexes. Alternatively, the dynamics of RNP transport in stressed cells could remain similar to those in basal conditions. In this scenario, the dissociation of ribosomes from mRNAs that accompanies stress-induced translational repression (Kedersha *et al*, 2000) could render transcripts more prone to forming RNA-RNA interactions when they are brought together by motors, thus promoting formation of stress granules (Parker *et al*, 2025; Van Treeck & Parker, 2018). Distinguishing between these possibilities through three-dimensional tracking of stress granule precursors is an important, albeit challenging, goal of future studies. It also remains to be determined whether FXR proteins, BICD2 and dynein have broader roles in RNP transport under basal conditions, including in the context of FXR1-containing RNP networks in cancer cell lines (Chen *et al*, 2024) or the establishment of polarized mRNA distributions in specialized cells such as neurons (Das *et al*, 2019).

## ACKNOWLEDGMENTS

We thank all members of the Bullock group, as well as many other colleagues in the LMB, for advice and support. We are particularly grateful to the following at the LMB: Florence Young (Bullock group), Giulia Manigrasso (Carter group), Steven Wingett (Lancaster group), Farida Begum, Mark Skehel and Catarina Franco (Mass Spectrometry Facility), Pier-Andrée Penttilä (Flow Cytometry Facility), Adrian Perrin, Elfy Chiang and Shraddha Nayak (VisLab), and Stephen McLaughlin (Biophysics Facility). We also thank Madison Edwards and Simpson Joseph (UC San Diego) for reagents and advice on recombinant FXR protein production. This work was supported by the Medical Research Council as part of UK Research and Innovation (UKRI) (intramural funding to S.L.B. (reference number MC_U105178790) and PhD studentships to Y.N.A.J. and E.M.), as well as an LMB-AstraZeneca Blue Sky award (to L.A.-A. and S.L.B.).

## DATA AND MATERIALS AVAILABILITY

Materials generated in this study, including engineered cell lines and plasmid constructs described in the Materials and Methods, are available from the corresponding author upon request.

## MATERIALS AND METHODS

### Plasmids

Sequences encoding the canonical isoforms of human DYNC1I2 (UniProt accession number Q13409-1), BICD2 (Q8TD16-1), FXR1 (P51114-1), FXR2 (P51116) and G3BP1 (Q13283-1) were cloned by Gibson assembly into the pcDNA5-FRT/TO-eGFP-TEV plasmid (provided by A. Castello [University of Glasgow, UK]) for tetracycline-inducible expression in human cells of proteins tagged at the N-terminus with GFP. For experiments assessing suppression of stress granule phenotypes in FXR-deficient cells by wild-type and Δ178-200 FXR1 transgenes, gene synthesis (Twist Bioscience) was used to introduce synonymous mutations into the full-length FXR1 coding sequences (aa 1–621) that render them resistant to the FXR1 siRNAs. pOG44 (Invitrogen, Thermo Fisher Scientific) was used to provide Flp recombinase for site-specific integration of pcDNA5-FRT/TO-eGFP-TEV-derived plasmids. pIDC-LIC2-IC2C-Tctex1-Robl1-LC8 and pACEBac1-His-ZZ-LTLTL-SNAP-DYNC1H1 (Schlager *et al*, 2014) were used to generate baculovirus for expression of the human dynein complex in Sf9 insect cells. Plasmids for expression of His- and MBP-tagged human FXR proteins in *E. coli* (isoform 2; Edwards *et al* 2020) were provided by M. Edwards and S. Joseph (UC San Diego, USA). These plasmids code for versions of the FXR proteins that lack ~90 amino acids from the disordered C-terminal tail that impair protein stability (Edwards *et al*, 2020). Plasmids for expression of Strep-tagged variants of mouse BICD2 (Q921C5) in Sf9 insect cells were provided by G. Manigrasso (Manigrasso *et al*, 2025; MRC-LMB, UK). These plasmids contain mouse BICD2 sequences that are codon optimized for expression in Sf9 insect cells. The full-length mouse and BICD2 proteins have an average amino acid identity of ~93%, with ~99% amino acid identity in CC3. The GST-BICD2^CC3^ plasmid for expression of mouse BICD2 sequences in *E. coli* was constructed in pGEX-4T-1 Truncations to coding sequences were generated by Gibson assembly of PCR-generated fragments. Mutations were introduced by oligonucleotide-based side-directed mutagenesis. Plasmid sequences were verified by whole-plasmid sequencing before use (Plasmidsaurus).

### Cell culture and cell line generation

Sf9 cells (Oxford Expression Technologies Ltd) were cultured at 27 °C in Insect-XPRESS protein-free medium containing L-Glutamine (Lonza). HeLa and U2OS cells were cultured at 37 °C with 5% CO_2_ in complete DMEM (high-glucose DMEM supplemented with 1 x GlutaMax [Gibco], 10% fetal bovine serum [Gibco] and 1% penicillin/streptomycin solution [Gibco]). Flp-In HeLa and U2OS cells were generated with the Flp-In™ T-Rex system (Invitrogen, Thermo Fisher Scientific) and cultured in complete DMEM supplemented with 150 μg/mL hygromycin and 5 μg/mL blasticidin (both from Gibco). Flp-In cell lines expressing GFP only were previously described (Madan *et al*, 2023). Populations of U2OS cells expressing GFP-FXR1 or GFP-FXR1^Δ178-200^ were sorted by flow cytometry to ensure similar levels of GFP expression. Where required, oxidative stress was induced by treatment with 300 μM sodium arsenite (NaAsO_2_; Sigma-Aldrich) in complete DMEM for the indicated time periods. At the onset of the project, cells were certified as Mycoplasma-free using the MycoAlert kit (Lonza).

### GFP nanobody immunoprecipitations

Expression of GFP-tagged proteins, or GFP alone, in HeLa Flp-In cell lines was induced for a minimum of two days prior to immunoprecipitations by supplementing the media with tetracycline hydrochloride (Gibco). 10 μg/mL tetracycline hydrochloride was used in all cases with the exception of the experiment that assessed interactions with the GFP-tagged variants of BICD2^CC3^, which were expressed at relatively low levels. To minimize differences in the expression of these constructs compared to the GFP-only control in these experiments, the concentration of tetracycline hydrochloride for the control cells was reduced to 0.5 μg/mL. Immunoprecipitations that were analyzed by immunoblotting typically used a single 15-cm dish of cells per sample, whereas those analyzed by mass spectrometry used two 15-cm dishes of cells per sample, with three samples processed per condition.

Cells were harvested with sterile cell scrapers after washing twice with ice-cold phosphate buffered saline (PBS). Following pelleting by centrifugation, cells from each 15-cm plate were lysed in 200 μL of buffer comprising 5 mM Tris-HCl pH 7.5, 120 mM NaCl, 0.25 mM EDTA, 1 mM phenylmethylsulfonyl fluoride (PMSF), 1X cOmplete EDTA-free Protease Inhibitor (Roche), 1X PhosSTOP (Sigma-Aldrich), and 0.5% NP-40 (Sigma-Aldrich) for 30 min on ice, with several vigorous passes through a pipette every 10 min. Following clearing by centrifugation at 20,000 *g* for 10 min at 4 °C, the lysates were transferred to a new tube and diluted with one volume of dilution buffer (5 mM Tris-HCl pH 7.5, 75 mM NaCl, 0.25 mM EDTA, 1X cOmplete EDTA-free Protease Inhibitor, 1X PhosSTOP phosphatase inhibitor) for each two volumes of lysate. Protein concentrations of lysates were measured with a Bradford assay kit (Thermo Fisher Scientific) and, if necessary, harmonized across samples by further dilutions. 500 μL of diluted lysate sample was incubated with 25 μl of pre-equilibrated magnetic agarose GFP-trap beads (Chromotek) in dilution buffer.

Sensitivity of interactions to the presence of RNA were assessed by addition of RNase A (Ambion, Life Technologies; final concentration 25 μg/mL) or RNasin RNase inhibitor (Promega; final concentration 0.5 U/μL) to the lysate just before the bead incubation. Tubes were then incubated for 1.5–2.5 h at 4 °C with end-over-end rotation. The beads were subsequently rinsed once with 500 μL ice-cold ‘Citomix’ wash buffer (60 mM KCl, 0.075 mM CaCl_2_, 5 mM K_2_HPO_4_/KH_2_PO_4_ pH 7.6, 12.5 mM 4-[2-hydroxyethyl]-1-piperazineethanesulfonic acid [HEPES] pH 7.6, 2.5 mM MgCl_2_, 0.1% [v/v] Triton X-100, 2 mM ATP, 1× cOmplete EDTA-free protease inhibitor, 1× PhosSTOP phosphatase inhibitor) before transfer to a fresh tube and a further rinse with 500 μL ice-cold Citomix wash buffer.

For experiments analyzed by immunoblotting, 40 μL 4× NuPAGE LDS sample buffer (Life Technologies) containing 200 mM dithiothreitol (DTT) was added to the beads followed by incubation at 98 °C for 15 min and storage of samples at −20 °C. For experiments analyzed by mass spectrometry, washed beads were subjected to two further brief washes with 50 mM ammonium bicarbonate (pH 8.0) before storage at −20 °C in a minimum volume of the solution.

### Label-free quantitative (LFQ) mass spectrometry

Proteins were digested on beads and processed for mass spectrometry as described previously (Madan *et al*, 2023). Data-dependent acquisition was performed in a Thermo QExactive Classic with an MS1 resolution of 35,000 over a mass-to-charge ratio range of 350– 1,600 (AGC target of 1 x 10^6^ and maximum injection time of 50 ms), followed by generation of ten separate tandem mass spectrometry spectra using higher-energy collisional dissociation at a normalized collision energy of 27, with 17,500 resolution, an AGC target of 2 x 10^5^ (with a maximum fill time of 100 ms), and a dynamic exclusion time of 30 s. Data were searched against the UniprotKB human protein database using the Andromeda (MaxQuant) search engine, with results processed and statistically evaluated using Perseus software (Tyanova *et al*, 2016). The protein tables were refined by exclusion of common contaminants and identifications from the reverse database. LFQ intensity values were normalized against the median intensity of each sample using only the peptides that had intensity values recorded across all samples. Only proteins detected in all three triplicate samples of at least one experimental group were evaluated further.

After log_2_ transformation, any missing values were imputed using values randomly drawn from a normal distribution calculated for each sample, as described previously (Neufeldt *et al*, 2019; Plaszczyca *et al*, 2019). Significant enrichment of proteins between conditions was evaluated by performing Welch’s t-tests with permutation-based false discovery statistics. Perseus was used to produce Volcano plots. STRING 11.0 (Szklarczyk *et al*, 2019) was used to display potential connectivity between enriched proteins using only experimental interaction data types and a medium-confidence (0.400) interaction score. Gene Ontology functional enrichment analysis was also performed in STRING, with proteins ranked based on false discovery rate (FDR).

### Electrophoresis and immunoblotting

Samples denatured in NuPAGE LDS buffer and DTT were electrophorized in precast 4–12% Bis-Tris NuPAGE gels (Life Technologies) in 1X 3-(N-morpholino)propanesulfonic acid (MOPS)-sodium dodecyl sulfate (SDS) buffer (Formedium) alongside Full-Range ECL™ Rainbow™ molecular weight markers (Sigma-Aldrich) or the Spectra Multicolor Broad Range protein ladder (Thermo Fisher Scientific). Proteins were wet-transferred onto Immobilon-P PVDF membrane (Millipore) using the X-cell SureLock system (Life Technologies). The membrane was then blocked in 5% (w/v) milk powder (Marvel) in PBS for 1 h at room temperature (RT) and cut horizontally to allow probing of multiple proteins per experiment. Following incubation with primary antibodies (Table S1) overnight at 4 °C or for 1 h at RT in 1% milk (w/v) in PBS, membranes were washed three times for 10 min in PBS at RT, incubated for 50 min with secondary antibodies (Table S1) in PBS at RT, and washed three times (10 min each) with PBS at RT. Signals were developed with the Enhanced Chemiluminescence (ECL) Prime Western Blotting Sytem (Cytiva) and visualized with Super RX-N medical X-ray film (FUJIFILM) and a JP-33 X-ray Film Processor (JPI Healthcare).

### Production of recombinant proteins

#### FXR1 and FXR2

FXR1 and FXR2, tagged with His-MBP, were expressed in *E. coli* and purified using a modified version of the protocol described in Edwards *et al* (2020). For each protein, clarified lysate from 2 L of Rosetta cells (Novagen) was treated with concentrated ammonium sulfate in 5 mM HEPES pH 7.5 for at least 1 h at 4 °C. Precipitated protein was pelleted and resuspended in 75 mL lysis buffer (50 mM Tris pH 7.5, 1 mM EDTA, 1 mM DTT, 1 mM PMSF) and dialyzed in 2 L of dialysis buffer (50 mM Tris pH 7.5, 1 mM EDTA, 1 mM DTT) overnight using 3.5 K MWCO SnakeSkin dialysis tubing (Thermo Fisher Scientific). The dialyzed solution was then centrifuged at 3,900 *g* for 30 min at 4°C to remove insoluble protein. The supernatant was loaded onto a 5-mL heparin column (HiTrap heparin HP 5 mL [GE Healthcare]) by fast protein liquid chromatography (FPLC; ÄKTApurifier [Cytiva]) and bound proteins eluted by increasing salt concentration (25 mM Tris pH 7.5, 1 mM EDTA, 1 mM DTT, 25% (v/v) glycerol, 0–0.5 M NaCl for 15 column volumes, 25 mM Tris pH 7.5, 1 mM EDTA, 1 mM DTT, 25% (v/v) glycerol, 0.7 M NaCl for 5 column volumes, and 25 mM Tris pH 7.5, 1 mM EDTA, 1 mM DTT, 25% (v/v) glycerol, 0.7–1 M NaCl for 5 column volumes). Peak fractions were pooled and concentrated by centrifugation through a 15-mL 30 kDa MW cutoff centrifugal filter (Amicon Ultra-15, Millipore, Sigma-Aldrich) at 3,900 *g* and 4 °C.

#### BICD2

Recombinant Strep-tagged BICD2 proteins were produced in *Sf9* insect cells. Pellets from 1 L of expression culture were resuspended in 45 mL lysis buffer (30 mM HEPES pH 7.2, 300 mM NaCl, 1mM DTT, 2mM PMSF) supplemented with 1 x cOmplete-EDTA protease-inhibitor and lysed using 20–25 strokes of a Dounce tissue homogenizer. Lysates were cleared by centrifugation at 35,000 *g* for 45 min at 4 °C, followed by filtering with a GF filter (Sartorius). Cleared lysates were then incubated with 1.5 mL of StrepTactin Sepharose beads (IBA Lifesciences) for 1 h at 4 °C on a rotator. Beads were washed on a gravity column with 200 mL wash buffer (30 mM HEPES pH 7.2, 300 mM NaCl, 1mM DTT), and proteins eluted with 5–7 mL of wash buffer supplemented with 3 mM desthiobiotin. Eluted proteins were concentrated to a volume of ~0.5 mL and loaded onto a Superose 6 Increase 10/300 column (Cytiva) that was pre-equilibrated in GF150 buffer (25 mM HEPES pH 7.5, 150 mM KCl, 1 mM MgCl_2_, 1 mM DTT). Peak fractions were pooled and concentrated to 1.5–6 mg/mL, depending on the construct.

GST and GST-BICD2^CC3^ were expressed in *E. coli* Rosetta 2 (DE3)pLysS cells. A 10 mL starter culture was used to inoculate 1 L of Luria–Bertani broth (LB) containing 25 μg/mL chloramphenicol and 100 μg/mL ampicillin, which was grown to OD600 0.4–0.6 and incubated with 0.6 mM IPTG at 14 °C for a further 16–18 h. Cells were harvested by centrifugation at 4500 *g* for 15 min at 4°C and the washed pellet resuspended in PBS containing 0.1% Triton X-100, 1 x Complete EDTA-free protease inhibitors, and 0.2 mM PMSF. All subsequent steps were performed at 4 °C or on ice. Cells were lysed by sonication and cleared by centrifugation at 50,000 *g* for 30 min. Proteins were purified by affinity chromatography using a GSTrap column (Cytiva) and eluted in PBS containing 0.1% Triton X-100, 1 x Complete EDTA-free protease inhibitors, 0.2 mM PMSF, and 10 mM glutathione. Protein-containing fractions were concentrated using Amicon Ultra centrifugal filters with a 3 kDa molecular weight cutoff (Millipore, Sigma-Aldrich).

#### Dynein and dynactin complexes

Recombinant human dynein complexes were purified from Sf9 cells using the protocol described by Schlager *et al* (2014) following infection with baculovirus that contained recombined sequences from pACEBac1-His-ZZ-LTLTL-SNAP-DYNC1H1 and pIDC-LIC2-IC2C-Tctex1-Robl1-LC8. This results in expression of dynein complexes containing: SNAP-tagged DYNC1H1, DYNC1LI2, DYNC1I2, Tctex1, Robl1 and LC8. Dynein complexes were captured via an interaction of the ZZ-tag on DYNC1H1 with an IgG Sepharose 6 affinity resin (Cytiva) within a gravity flow Econo-column (Bio-Rad) whilst labeling the SNAP domain with SNAP-Cell TMR-Star (New England Biolabs). Following elution with TEV protease, dynein complexes were further purified by FPLC. Native dynactin complexes were purified from pig brain, as described previously (Schlager *et al*, 2014; Urnavicius *et al*, 2015).

Aliquots of each of the protein samples were flash-frozen in liquid nitrogen and stored at −80 °C. The purity of each preparation was subsequently confirmed by SDS-PAGE.

### Differential scanning fluorimetry

Thermal denaturation of FXR1 proteins was followed using intrinsic protein fluorescence measured with a Prometheus NT48 instrument (Nanotemper Technologies). Samples were loaded into standard capillaries at a final concentration of 5.5 μM for wild-type FXR1 or 8 μM for FXR1^Δ178-200^ in a buffer comprising 25 mM Tris pH 7.5, 1 mM EDTA, 1 mM DTT, 25% glycerol, 0.7 M NaCl and heated at 2 °C/min from 15 to 95 °C. The first derivatives of 350/330 nm fluorescence emission ratios were analyzed using PR.ThermoControl software (v.2.3.1; Nanotemper Technologies).

### Pulldowns with purified proteins

In the absence of RNA targets, FXR1 and FXR2 are unstable in physiological salt concentration concentrations, presumably due to imbalanced charge distribution (Edwards *et al* 2020). To overcome this issue, the binding steps of pulldowns involving these proteins were performed in buffer containing a high salt concentration (600 mM NaCl) (Edwards *et al* 2020).

For StrepTag-based pulldowns, 20 μL MagStrep Type3 XT bead slurry (IBA Lifesciences) was equilibrated in pre-cooled dilution buffer (5 mM Tris-HCl pH 7.5, 600 mM NaCl, 0.25 mM EDTA, 1X cOmplete EDTA-free Protease Inhibitor [Roche], 1X PhosSTOP [Sigma-Aldrich]), followed by incubation with 2.85 μg of Strep-tagged full-length or truncated mouse BICD2 (provided by G. Manigrasso, MRC-LMB [Manigrasso *et al*, 2025] or expressed and purified as described above). To assess interactions of FXR proteins with BICD2, 5 μg of His-MBP-FXR1 or His-MBP-FXR2 were combined with 500 μL of dilution buffer and added to the beads. The tubes were then subjected to end-over-end rotation for 1.5–2 h at 4 °C, followed by washing of beads six times with 500 μL ice-cold wash buffer (600 mM NaCl, 5 mM MgCl_2_, 5 mM Tris HCL, 1x PhosSTOP, 1X cOmplete EDTA-free Protease inhibitor, 0.1% Triton X-100). For the final wash, the beads were transferred to fresh Eppendorf tubes. Samples were processed for immunoblotting as described above.

For GST-based pulldowns, the protocol above was followed with the following exceptions: 20 μL Glutathione Magnetic Agarose Beads (Pierce) were used; the dilution buffer did not include EDTA; 2.65 μg GST-BICD2^CC3^ and 1.38 μg of GST were used (0.1 μM final concentration of each protein) in combination with 0.2 μM FXR1 in the binding step; the wash buffer was the dilution buffer containing 0.1% Triton X-100.

To assess the association of dynein with FXR proteins, recombinant human dynein was coupled to Protein G magnetic beads (Pierce) via antibodies to DYNC1I2 (Sigma-Aldrich, MAB1618), according to the manufacturer’s instructions. In control experiments, beads were coupled to FLAG antibodies (Sigma-Aldrich, F3165). Binding assays were performed as described above for the StrepTag-based pulldowns.

### siRNA transfection

SMARTpools of four ON-TARGETplus siRNAs were synthesized by Dharmacon and resuspended in 1X siRNA buffer (Dharmacon) to a stock concentration of 20 μM. The siRNAs used targeted the following genes: BICD1 (L-019496-00-0010), BICD2 (L-014060-00-0010), DYNC1H1 (L-006828-00-0005), FXR1 (L-012011-00-0020) and FXR2 (L-011955-00-0020). These pools cause strong reductions in expression of the target genes (see Figure S6A for validation of knockdown for siRNAs targeting FXR and BICD proteins, as well as Madan *et al* (2023) and Albacete-Albacete *et al* (2026) for validation of siRNAs targeting DYNC1H1). A non-targeting SMARTpool (ON-TARGETplus D-001810-10-20; siCTRL) was used as a negative control. For knockdowns coupled to immunoprecipitations, 3 x 10^6^ cells were seeded in 15-cm dishes one day before siRNA transfection. 69 μL Lipofectamine RNAiMax reagent (Invitrogen), 3.8 mL OptiMEM (Gibco) and 38 μL of siRNA stock solution (or 19 μL for each SMARTpool when knocking down paralogs) were pre-incubated for 5 min at RT before addition to 20 mL of culture medium. Cells were harvested 72 h after transfection and processed for immunoprecipitation, as described above. For knockdown experiments analyzed by light microscopy, cells were seeded in 24-well plates (for analysis of fixed cells) or 8-well glass-bottomed Ibidi μ-Slides (Thistle Scientific; for analysis of live cells) one day prior to siRNA introduction (for forward transfection) or on the same day (for reverse transfection), with the volumes of the transfection reagents and siRNAs adjusted accordingly. To monitor stress granule formation, media containing arsenite, or control media, was used to replace the media 72 h after siRNA treatment.

### Analysis of RNA content of immunoprecipitates

Immunoprecipitations were performed from HeLa cell extracts as described above, except that Citomix buffer was replaced with a buffer comprising 60 mM NaCl, 5 mM MgCl_2_, 5 mM Tris-HCl, 1X cOmplete EDTA-free protease inhibitor, 1x PhosSTOP phosphatase inhibitor and 0.1% Triton X-100. For RNA isolation from these samples, washed protein-bound beads were incubated with TRIzol™ reagent (Thermo Fisher Scientific) for 5 min at RT, followed by addition of 60 µL chloroform. Samples were gently mixed for 2–3 minutes and centrifuged at 12,000 *g* for 15 min at 4°C. The aqueous phase was collected without disturbing the interface and the RNA content purified with the RNeasy MinElute kit (QIAGEN). RNA was eluted in 14 µL nuclease-free water and stored at −80 °C if not used immediately. RNA concentration and integrity were assessed using the Qubit™ RNA HS Assay Kit (Thermo Fisher Scientific) and RNA 6000 Pico Chips on a Bioanalyzer (Agilent). All samples had an ‘RNA integrity number’ >9, confirming suitability for downstream analysis.

cDNA synthesis from purified RNA samples was performed using the AffinityScript qPCR cDNA Synthesis Kit (Agilent) with random primers. cDNA synthesis was carried out at 42 °C for 30–45 min. Quantitative PCR (qPCR) reactions were prepared with SYBR Green master mix (Applied Biosystems), 0.5 pg/µL cDNA from the immunoprecipitated material, 0.5 pg/µL exogenous spike-in cDNA (human *XCR1*, which is not expressed in HeLa cells [Dorner *et al*, 2009]) and 0.3 µM primers targeting exon-exon junctions (Table S4) to ensure specificity for mature mRNA transcripts. Each reaction was performed in triplicate per sample using 10 μL total reaction volumes in 96-well plates. Plates were sealed with MicroAmp optical adhesive film (Applied Biosystems), spun at 1,000 rpm for 1 min at RT in an Eppendorf 5810 centrifuge, and run on a ViiA™ 7 Real-Time PCR System (Applied Biosystems) qPCR machine using cycling conditions recommended by the manufacturer. RNA abundance was normalized to that of the spike-in cDNA.

### RNA sequencing

RNA was extracted using the RNeasy extraction kit (QIAGEN), according to the manufacturer’s protocol. Total RNA concentration and RNA integrity were confirmed as described above. mRNA isolation was performed from 1 μg of total RNA per sample using the NEBNext poly(A) mRNA magnetic isolation module (NEB). Libraries were generated from mRNA samples with the NEB-Next Ultra II directional RNA library prep kit for Illumina (NEB) following the manufacturer’s instructions, with barcoding using Multiplex Oligos for Illumina (Dual index primers pairs set 4, NEB). Library concentrations, quality and average size were determined with Qubit dsDNA High Sensitivity (Invitrogen) and High Sensitivity DNA kits (Agilent). 780 pM of pooled barcoded libraries were sequenced with the NextSeq 2000 system (Illumina, NextSeq 2000 P3; 100 cycles in paired-end 100 mode at a minimum depth of 25 million reads per library).

RNAseq data were processed as follows. FASTQ files were subjected to quality control, trimming and mapping with the *rnaseq* pipeline (version 3.12.0) distributed by *nf-core* (https://zenodo.org/records/7998767). Each component within the pipeline was version controlled and executed within a Singularity container (https://sylabs.io). The pipeline was executed with the following command:

*nextflow run nf-core/rnaseq -r 3.12.0 --input sample-sheet.csv --genome homo_sapiens.GRCh38.release_102 -config lmb.config --outdir results --deseq2_vst -bg*

The parameters largely followed the default settings of the *nf-core rnaseq* pipeline, with STAR (version 2.7.9a) (Dobin *et al*, 2013) used for alignment and reads mapped to the human reference genome (GRCh38/release 102). To account for the Illumina color chemistry, the *--nextseq 20* read-trimming parameter was applied. A sample list was also provided, which specified forward and reverse FASTQ file pairings, sample identities, and replicate grouping. The *nf-core rnaseq* pipeline generated quantified expression values for each gene in each sample, both as raw counts and transcripts per million (TPM), which were used for downstream analyses. Differential gene expression analysis was performed with the DESeq2 statistical framework in SeqMonk (version 1.48.1), with raw read counts per gene used as input.

### *In vitro* reconstitution of transport complexes

Biotinylated porcine microtubules stabilized with GMP-CPP (guanosine-5’-[(α,β)-methyleno]triphosphate) were adhered to biotinylated glass coverslips within imaging chambers via a streptavidin linkage using previously described protocols (McClintock *et al*, 2018). FXR proteins (or FXR buffer alone) were pre-incubated with mouse BICD2_FL_ in motility buffer (30 mM HEPES, 5 mM MgSO_4_, 1 mM ethyleneglycoltetraacetic acid (EGTA), 1 mM DTT) on ice for 20 min before the addition of dynein and dynactin and a further incubation of 40–100 min on ice. In control experiments, BICD2 was omitted. The molar ratios of dynein:dynactin:BICD2_FL_: FXR1/FXR2 in the assembly mix were 1:2:5:9.5/11 (704 nM dynein, 1.408 μM dynactin, 3.52 μM BICD2_FL_, 8.25 μM FXR1/9.45 μM FXR2). After incorporating all the factors into the mix, the salt concentration was 100 mM. Just prior to imaging, the assembly mixes were diluted 40-fold in ‘dilution buffer’ to achieve a final buffer composition of 25 mM KCl, 1 mg/mL α-casein, 5 mM MgATP and an oxygen scavenging system comprising 1.25 μM glucose oxidase, 140 nM catalase, 71 mM 2-mercaptoethanol, and 25 mM glucose to minimize photobleaching during image acquisition. The diluted protein samples were immediately loaded into imaging chambers.

Imaging of fluorescent dynein in chambers was carried out at RT using a Nikon TIRF microscope system under the control of Micro-Manager (Edelstein *et al*, 2010) and equipped with a 100x/1.49-NA oil objective (Nikon APO TIRF) and Coherent Sapphire 488-nm (150 mW), Coherent Sapphire 561-nm (150 mW), and Coherent CUBE 641-nm (100 mW) lasers. Images were captured with an iXonEM+ DU-897E EMCCD camera (Andor), providing pixel dimensions of 105 x 105 nm. Images were captured at 2 frames/s and 100 ms exposure per frame. For experimental series in which BICD2 was excluded, dynein complexes were imaged with a 100x/1.49-NA APO TIRF oil objective (Nikon) using the 561-nm laser within the Multiline Kompact laser box (Cairn) as part of a Nikon TIRF system controlled by Micro-Manager and fitted with the iLas2 platform for RING-TIRF illumination (GATACA Systems) at 200 ms per frame. Images on this microscope were acquired on a Photometrics Prime 95B CMOS camera at 2 frames/s, providing pixel dimensions of 108 x 108 nm.

### Analysis of *in vitro* motility data

Kymographs of *in vitro* motility data were generated and analyzed manually using Fiji, as described (McClintock *et al*, 2018). The position of microtubules was determined using either the fluorescent tubulin signal or a maximum projection of the dynein signal throughout the duration of the movie. Typically, a set of 15 microtubules was selected for analysis across two movies for each chamber, based on their length and separation from other microtubules, before visualizing the motile properties of associated motor complexes. Subsequent quantification was performed whilst being blinded to sample identities using the Blind Analysis Tool in Fiji. Run velocities were calculated by measuring the total displacement of each continuous dynein movement and dividing the value by the duration of the event.

### Immunofluorescence

Cells were plated on circular No.1.5 13-mm coverslips (VWR) in 24-well plates and fixed with 4% paraformaldehyde (Sigma-Aldrich) in PBS for 15 min at 37 °C. When imaging cell lines with GFP-tagged components, expression was induced for at least two days prior to fixation by addition of tetracycline hydrochloride (see above). Fixed cells were washed twice with 1 mL PBS and permeabilized with 0.1% Triton X-100 in PBS for 5 min at 4 °C. Non-specific binding sites were blocked by incubating with 2% bovine serum albumin (BSA; Merck) in PBS for 1 h at RT before washing once in PBS and incubating with primary antibodies (Table S1) in 0.1% BSA in PBS for 1.5 h at RT. After washing by submerging five times in a beaker containing 0.1% BSA in PBS, the samples were incubated with a PBS solution containing DAPI (Sigma-Aldrich; final concentration 1 μg/mL) and secondary antibodies (Table S1) in 0.1% BSA for 1 h at RT in the dark. Samples were then washed in 0.1% BSA in PBS, followed by two washes in PBS, and mounting in Prolong Diamond Antifade Mountant (Life Technologies) on 76-mm x 26-mm microscopy slides (Thermo Fisher Scientific).

Imaging was performed either with (1) a laser scanning confocal Zeiss 710 UV inverted microscope equipped with a 63x/1.4-NA oil objective or (2) a laser scanning confocal Zeiss 780 UV inverted microscope equipped with a 40x/1.4-NA oil objective. Laser power and acquisition parameters were kept constant within a single experimental series.

### Proximity ligation assay

Close apposition between proteins in cells was visualized with the Duolink Proximity Ligation Assay (Sigma-Aldrich). The assay was performed following the manufacturer’s protocol, with probes to anti-mouse antibodies (detecting GFP) and anti-rabbit antibodies (detecting FXR1 or FXR2) (Table S1) coupled to the Duolink *InSitu* FarRed Detection Reagent. Imaging was performed as described in the ‘Immunofluorescence’ section. Acquired images were analyzed using ImageJ (Schneider *et al*, 2012). The number of PLA spots per cell was analyzed by creating a mask with the threshold tool and using the watershed algorithm to separate touching objects. Numbers of spots from negative controls, in which no primary antibodies were added, were subtracted from the spot number obtained in the experimental conditions.

### Live imaging of stress granule formation in GFP-G3BP1 U2OS cells

Cells were plated in 8-well glass-bottomed Ibidi μ-slides. Movies were acquired with a Nikon X1 Eclipse Ti Spinning Disk microscope using a 60x/1.2-NA water objective at a rate of 1 frame/15 s. The imaging system was equipped with a Photometrics Prime 95B sCMOS (95% QE) camera and a stage-top environmental control chamber (Okolab), which was set to 37 °C and contained 5% CO_2_. 1 h-long movies of stress granule formation were analyzed manually in Fiji by segmenting each cell, setting a normalized intensity threshold, and creating a mask of the GFP signal. Watershedding was used to further segment any touching stress granules, followed by quantification of the number of stress granules across the image series.

### Quantification of stress granule assembly in fixed cells

Stress granule phenotypes in U2OS cells stained with antibodies to G3BP1 (Table S1) were quantified using a script that was custom-produced in Python (https://github.com/jboulanger/sganalysis.git). Cell and cytoplasm segmentation was performed using a pre-trained network, Cellpose 2.0 (Pachitariu & Stringer, 2022), based on signals from Alexa555-Wheat Germ Agglutinin (which was incubated with the cells at a final concentration of 1 μg/mL prior to washing) and DAPI (used as described above). For experiments assessing the ability of wild-type and Δ178-200 GFP-FXR1 transgenes to suppress stress granule phenotypes in FXR-deficient cells, only cells with a mean GFP intensity that was at least four times higher than the mean fluorescence values for the unmodified U2OS cells in the same experiment were analyzed. This ensured that non-expressing cells in the GFP-FXR1 cell populations were not evaluated. To account for any differences in imaging sensitivity between independent experiments, GFP intensity values were normalized to this threshold.

