## Supplementary materials for "FXR proteins and BICD2 cooperate to promote dynein-mediated RNP transport and stress granule assembly"

**Supplementary Figures, Movie Legend and Supplementary Tables**

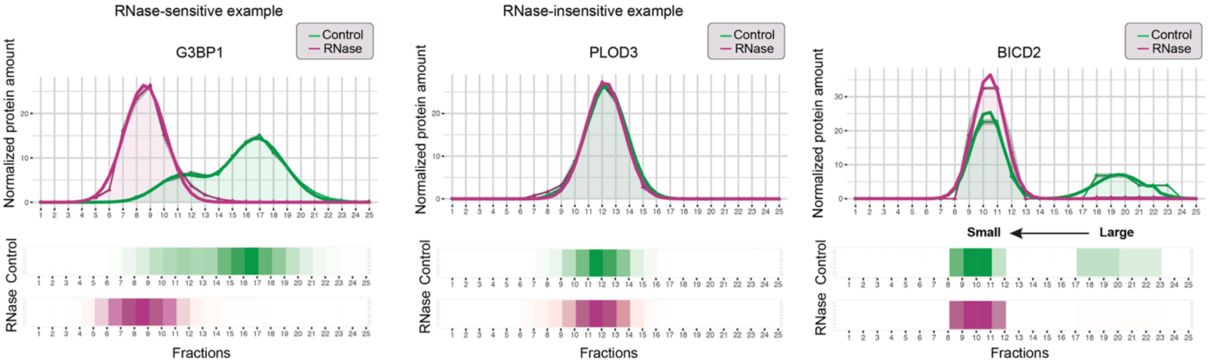


**Figure S1: Evidence that BICD2 is complexed with RNA.** Relative abundance of the indicated proteins in fractions collected after density gradient ultracentrifugation of HeLa cell lysates without (control) or with RNase treatment. Data are reproduced from the R-DeeP database (Caudron-Herger *et al*, 2019). G3BP1 and PLOD3 are examples of proteins with RNase-sensitive and RNase-insensitive profiles, respectively. A fraction of BICD2 is present in a relatively large, RNase-sensitive complex.

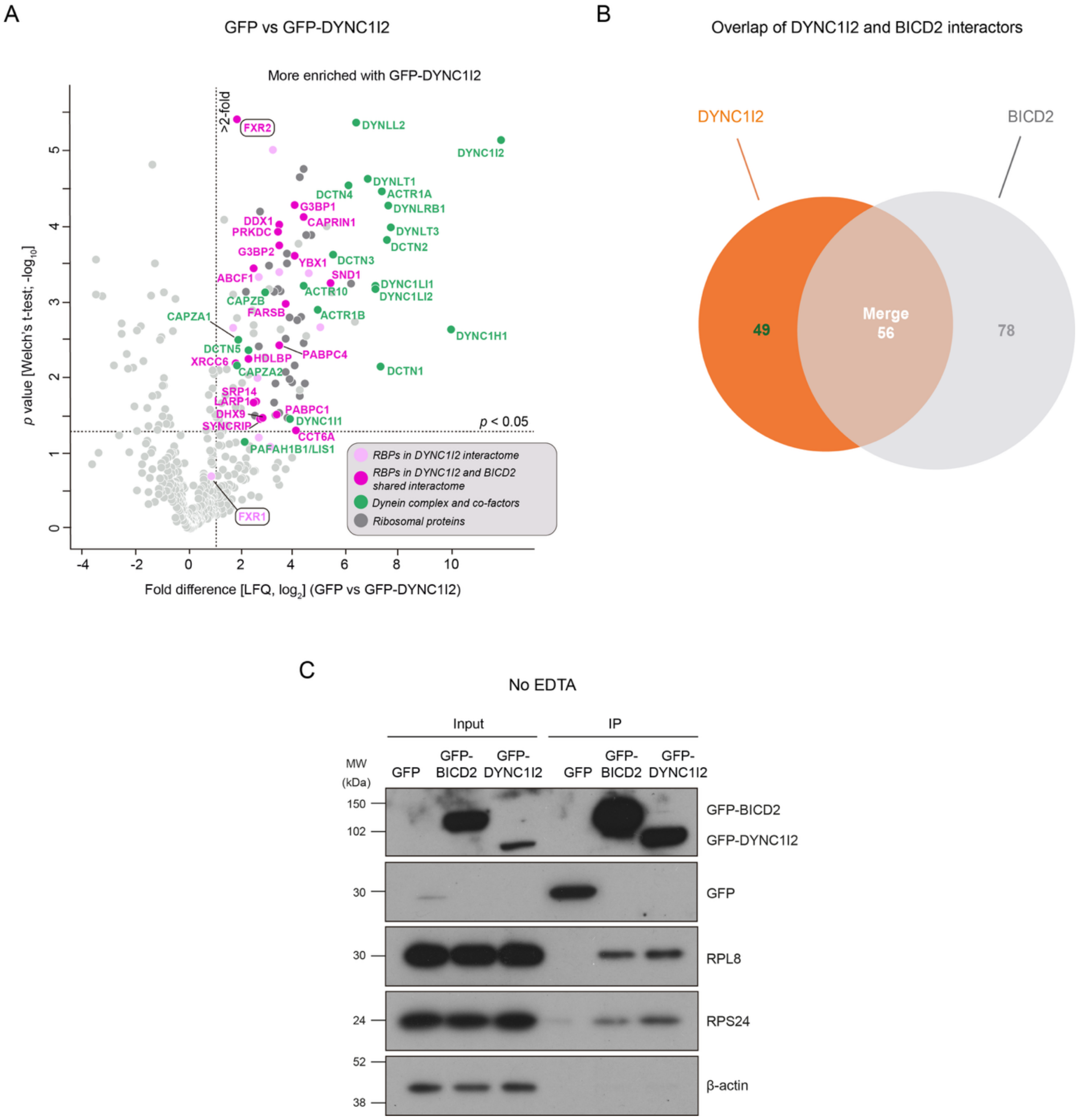


**Figure S2: Supplementary data from nanobody capture experiments. (A)** Volcano plot of mass spectrometry data showing enrichment of proteins and *p* values in GFP-DYNC1I2 vs GFP immunoprecipitates from HeLa cells (three biological replicates per condition). LFQ: label-free quantification. ACTR1A, ACTR1B, ACTR10, CAPZA1, CAPZA2, CAPZB and DCTN1–5 are components of the dynactin complex. **(B)** Venn diagram showing number of proteins significantly enriched (> 2-fold enrichment and $p < 0.05$) in GFP-DYNC1I2 and/or GFP-BICD2 immunoprecipitates relative to their respective GFP only controls. **(C)** Results of immunoblotting assessing the association of GFP-BICD2 with the large and small ribosomal subunits in HeLa cell lysates when ribosome splitting is prevented by omission of EDTA from the assay buffer. RPL8 and RPS24 are components of the large and small ribosomal subunits, respectively. MW: molecular weight markers. IP: immunoprecipitated material. β-actin serves as a specificity control for the immunoprecipitation.

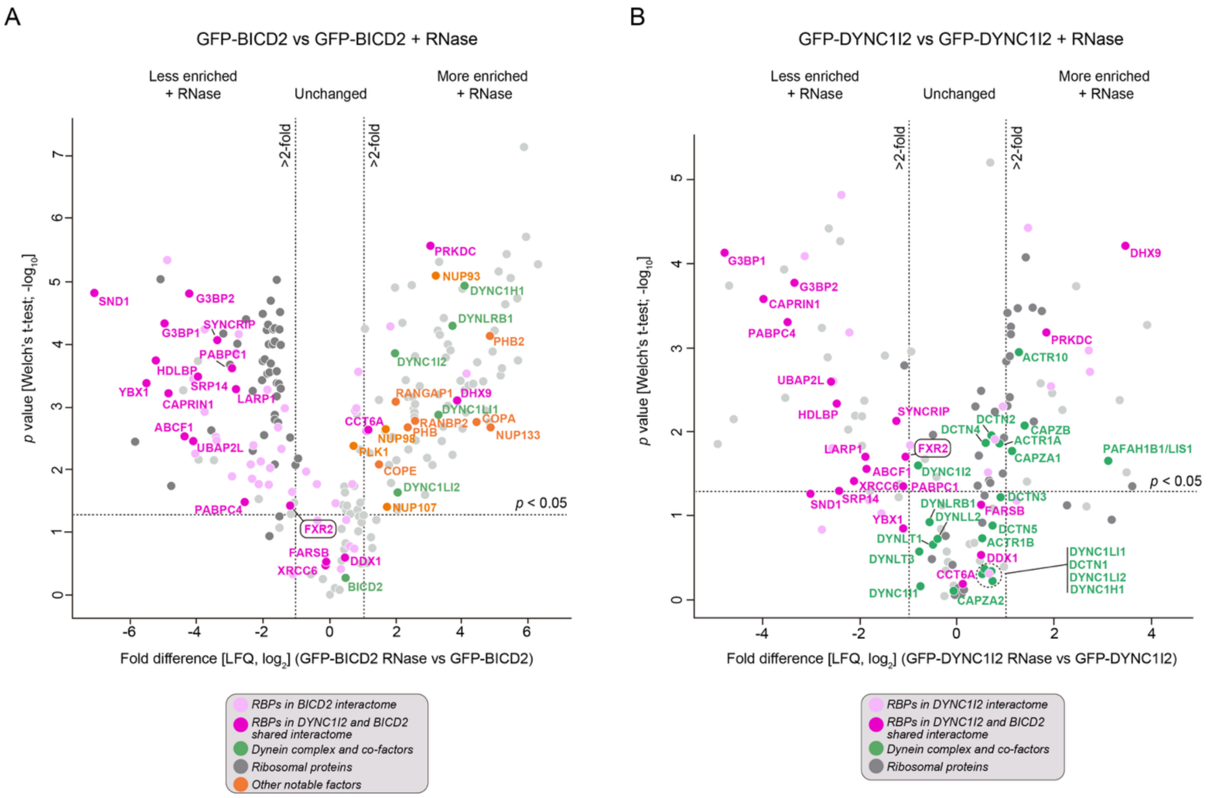


**Figure S3: Supplementary data on RNase sensitivity of BICD2 and DYNC1I2 interactions. (A, B)** Volcano plots showing enrichment of proteins in 'GFP-BICD2 with RNase' vs 'GFP-BICD2 without RNase' (A) and 'GFP-DYNC12 with RNase' vs 'GFP-DYNC1I2 without RNase' (B) immunoprecipitates from HeLa cells, as well as associated *p* values (three biological replicates per condition). The association of FXR2 with BICD2 and dynein in these experiments was not strongly dependent on RNA, as levels of FXR2 in the immunoprecipitates were reduced by only ~50% by RNase treatment. Note that RNase increases the association of BICD2 with several membrane-associated proteins (orange), raising the possibility of competition for BICD2 binding between these factors and RNase-sensitive RBP interactors. BICD2's interaction with PLK1 was not significantly affected by RNase, whereas its interaction with dynein components was increased by RNase. This could conceivably reflect the membrane-associated proteins whose interaction with BICD2 is increased by RNase strongly promoting recruitment of the motor complex.

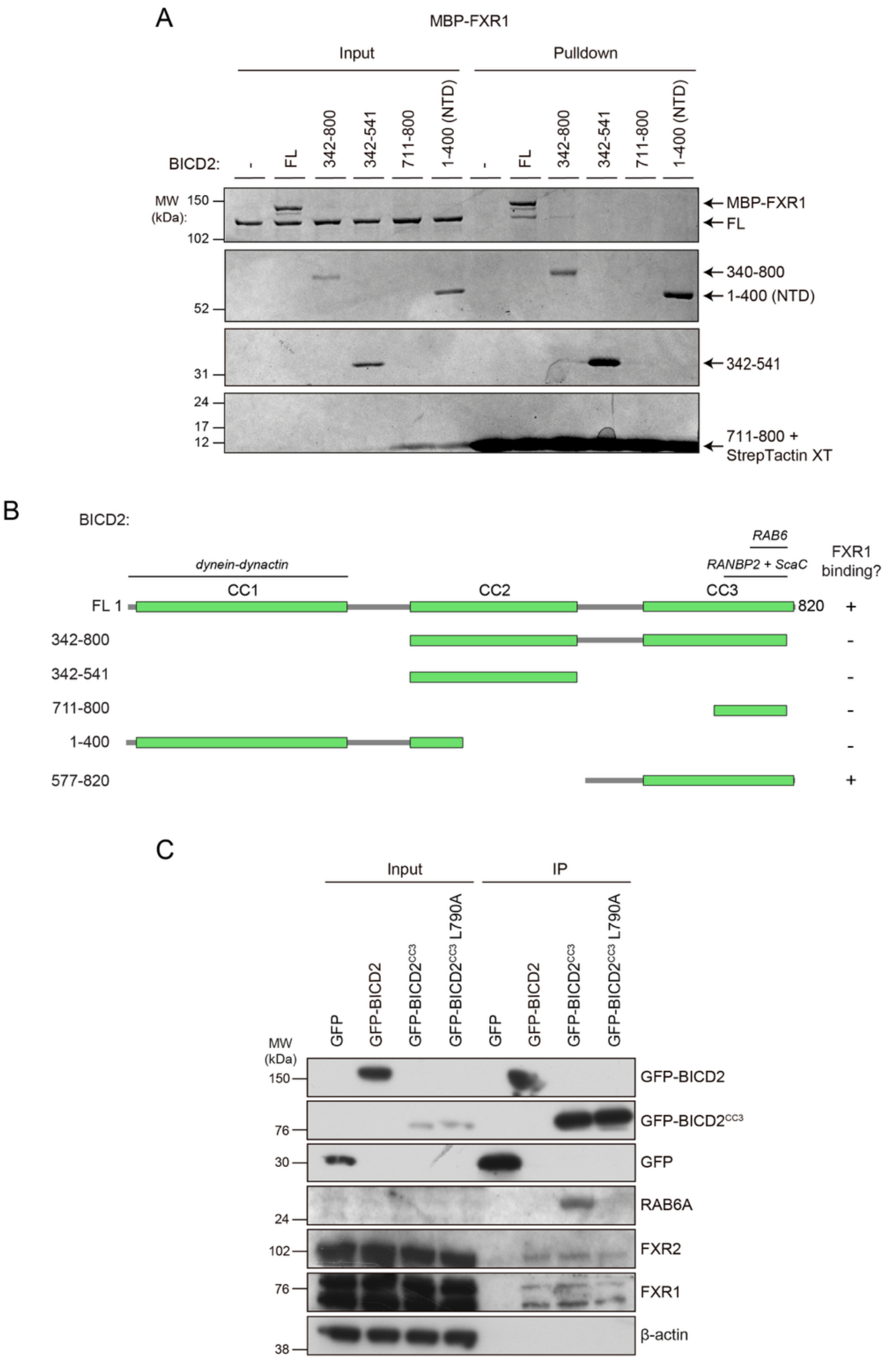


**Figure S4: Supplementary data on biochemical interactions of FXR proteins and BICD2. (A)** Images of Coomassie-stained gel showing results of *in vitro* pulldown assays with the indicated BICD2 truncations and MBP-FXR1. BICD2 proteins were immobilized on beads via a Strep-tag. Only $BICD2_{FL}$ captures FXR1. $BICD2^{711\text{-}800}$ cannot be distinguished from the Strep-tag-binding StrepTactin XT monomers, which have a similar molecular weight. **(B)** Summary of BICD2 variants used in the *in vitro* pulldown experiments and their ability to bind FXR1. Previously reported binding sites for known BICD2 partners are shown. Folded domains are shown in green. Note that CC3 ends at residue 814 (Parcoil2 prediction) and therefore is truncated in the 342–800 and 711–800 variants. **(C)** Immunoblots showing results of co-immunoprecipitation experiments assessing the ability of wild-type and L790A human $BICD2^{CC3}$ to associate with FXR1 and FXR2 in HeLa cells. RAB6A is shown as a positive control for the effect of L790A; note that RAB6A binds more effectively to $BICD^{CC3}$ than full-length BICD2 as its binding site is exposed by removal of the N-terminal domain (Hoogenraad *et al*, 2003). GFP-$BICD^{CC3}$ contains aa 587–823 of human BICD2. β-actin serves as a specificity control for the immunoprecipitations. MW, molecular weight markers.

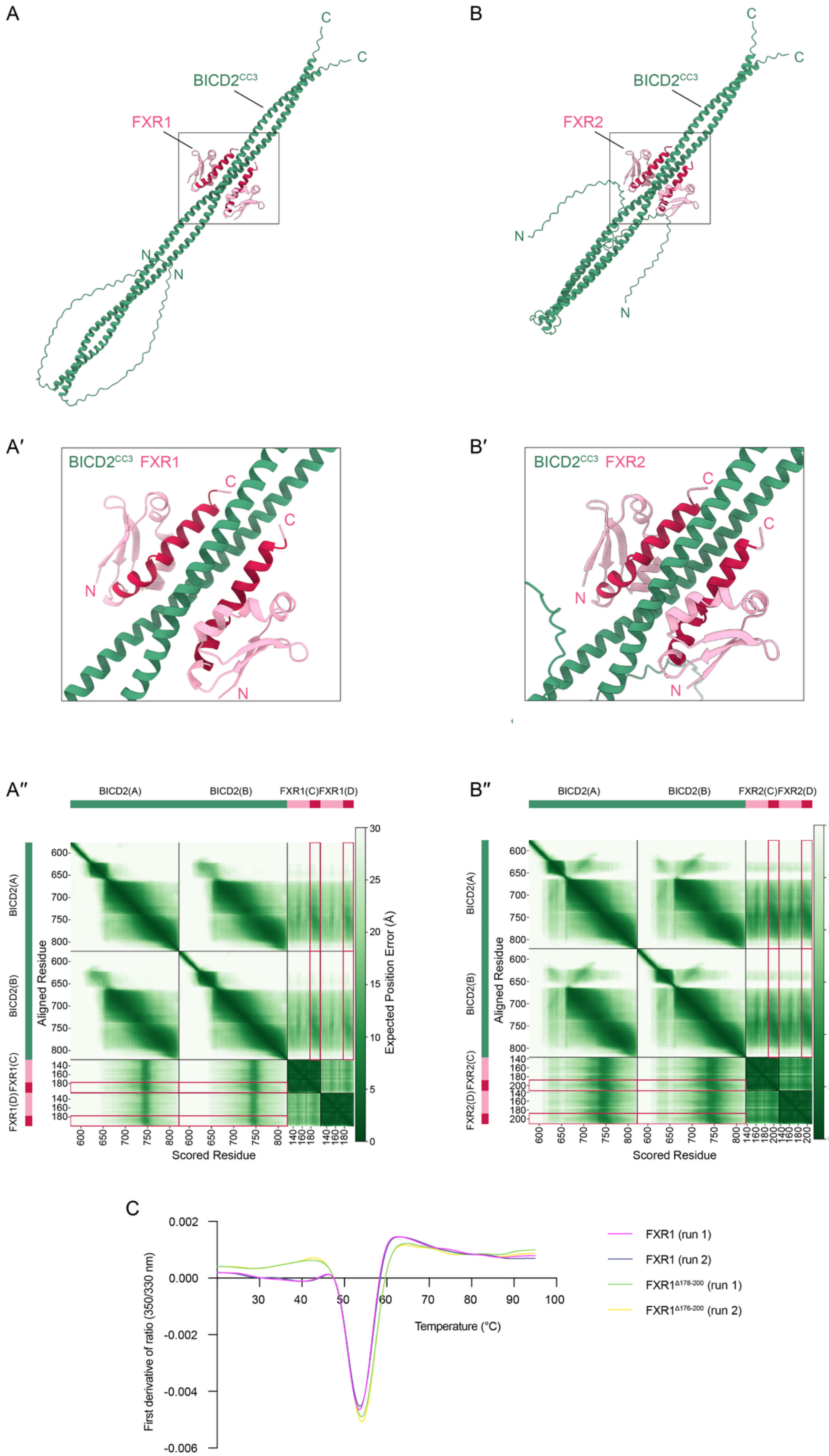
A
BICD2^CC3
FXR1
C
C
N
N
B
BICD2^CC3
FXR2
C
C
N
N
A′
BICD2^CC3 FXR1
B′
BICD2^CC3 FXR2
A″
BICD2(A)
BICD2(B)
FXR1(C)
FXR1(D)
Aligned Residue
Scored Residue
Expected Position Error (Å)
B″
BICD2(A)
BICD2(B)
FXR2(C)
FXR2(D)
Aligned Residue
Scored Residue
Expected Position Error (Å)
C
First derivative of ratio (350/330 nm)
Temperature (°C)
FXR1 (run 1)
FXR1 (run 2)
FXR1Δ178-200 (run 1)
FXR1Δ176-200 (run 2)

**Figure S5: Supplementary data on interaction of the FXR1 and FXR2 KH0 domains with BICD2 CC3.** **(A** and **B)** Highest confidence Alphafold3 predictions obtained using two BICD2 chains corresponding to the CC3 region and two KH0 domains of FXR1 (A) or FXR2 (B). Stoichiometry was based on that of other BicD-partner complexes. The same regions of FXR proteins (FXR1, 176–201; FXR2, 189–211) and BICD2 (734–761) interacted with each other in the top five predictions involving each FXR protein. A′ and B′ show magnified views of boxed regions in A and B, respectively. A″ and B″ show Predicted Aligned Error (PAE) plots for each complex. In each sub-panel, regions of FXR proteins that are deleted in the Δ178–200 FXR1 mutant, or the equivalent region of FXR2, are highlighted in crimson. In B, the N-terminal region of CC3 is folded back onto more C-terminal regions. **(C)** Differential scanning fluorimetry traces for purified wild-type and Δ178-200 FXR1 proteins. Both proteins have similar thermal denaturation profiles, indicating that the mutant protein is not misfolded.

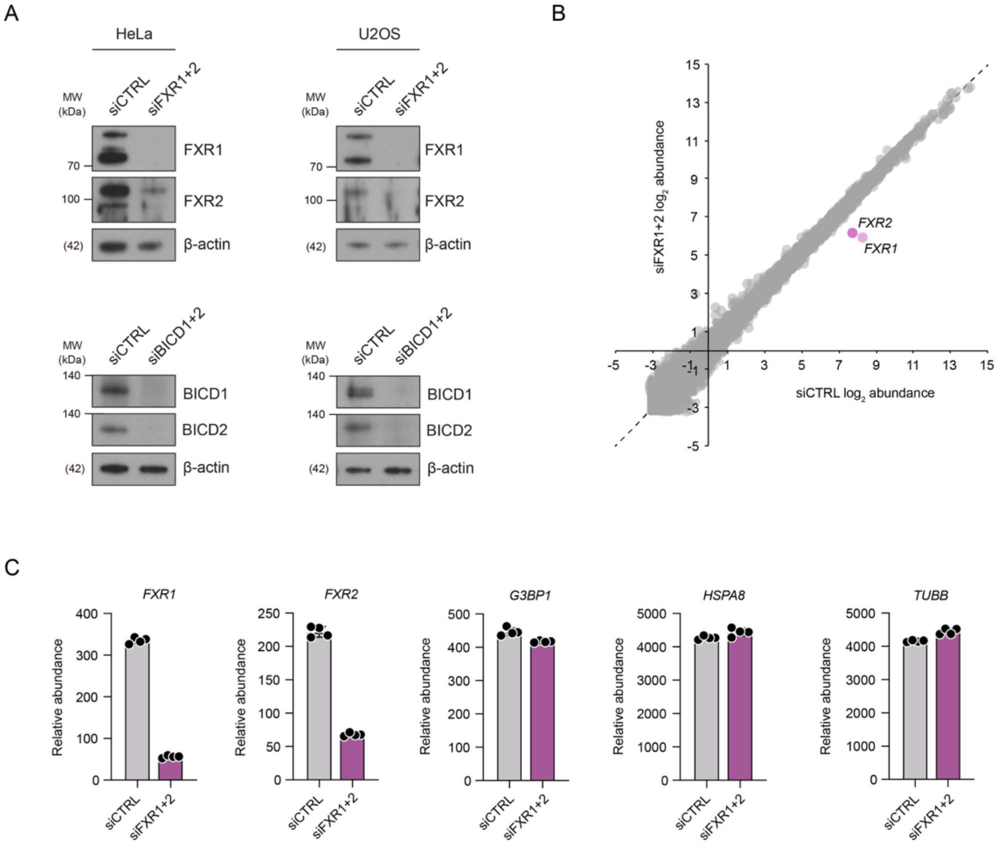


**Figure S6: Supplementary data on effects of siRNAs. (A)** Immunoblots showing efficient siRNA-mediated depletion of indicated target proteins in HeLa and U2OS cells. MW, molecular weight markers. In regions of the gel where there were no molecular weight markers, actual molecular weights of the protein of interest are shown in parentheses. **(B)** Scatterplot comparing relative mRNA abundance (mean $log_2$ normalized values from four biological replicates) determined by RNAseq in extracts of HeLa cells treated with siCTRL or siFXR1+2 RNAi pools. Only *FXR1* and *FXR2* mRNAs are substantially depleted in the siFXR1+2 cells (shrunk $log_2$ fold change > 1 and FDR < 0.05). **(C)** Relative abundance of the indicated FXR1 target mRNAs, as well as *FXR1* and *FXR2*, in the data presented in panel B.

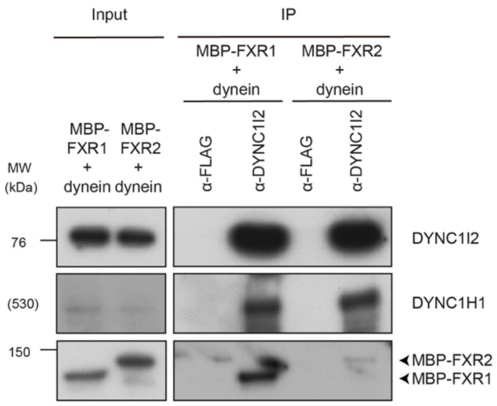


**Figure S7: Recombinant FXR1 and FXR2 can associate with dynein *in vitro*.** Immunoblots showing material captured using an α-DYNC1I2 or control α-FLAG antibody when MBP-FXR1 or MBP-FXR2 is incubated with the purified human dynein complex. Both MBP-FXR1 and MBP-FXR2 are captured with the dynein complex. IP, immunoprecipitation. MW, molecular weight markers. In regions of the gel where there were no molecular weight markers, actual molecular weights of the protein of interest are shown in parentheses.

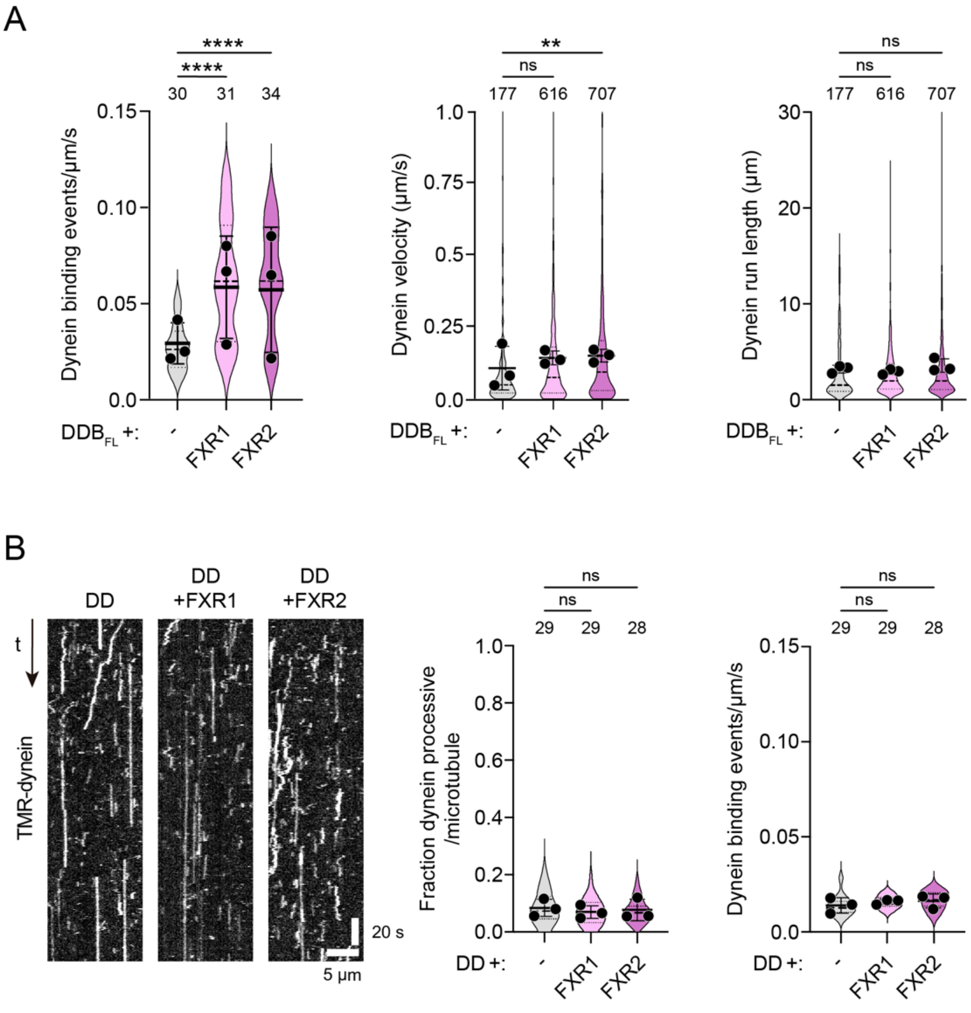


**Figure S8: Supplementary data for *in vitro* motility assays. (A)** Quantification of microtubule binding, velocity and run length of TMR-dynein in the presence of dynactin and full-length BICD ($DDB_{FL}$) when FXR1, FXR2 or buffer alone (-) are added. **(B)** Example kymographs (left) and quantification (right) of TMR-dynein behavior in the presence of dynactin and absence of BICD2 (DD) when FXR1, FXR2 or buffer alone (-) are added; t, time. In the absence of BICD2, neither FXR1 nor FXR2 stimulate dynein movement or microtubule binding. There were insufficient transport events in these conditions for meaningful quantification of run lengths and velocities. In A and B, violin plots represent data for individual microtubules, with circles representing mean values from individual experiments (N = 3 per condition). Solid lines and dashed lines within violins show median values and interquartile range, respectively. Numbers of microtubules are shown above bars. Statistical significance compared to $DDB_{FL}$ (A) or DD (B) was evaluated with a one-way ANOVA test with Tukey's post-hoc multiple comparisons correction using per microtubule values aggregated across experiments (ns, not significant; **, $p < 0.01$; ****, $p < 0.0001$).

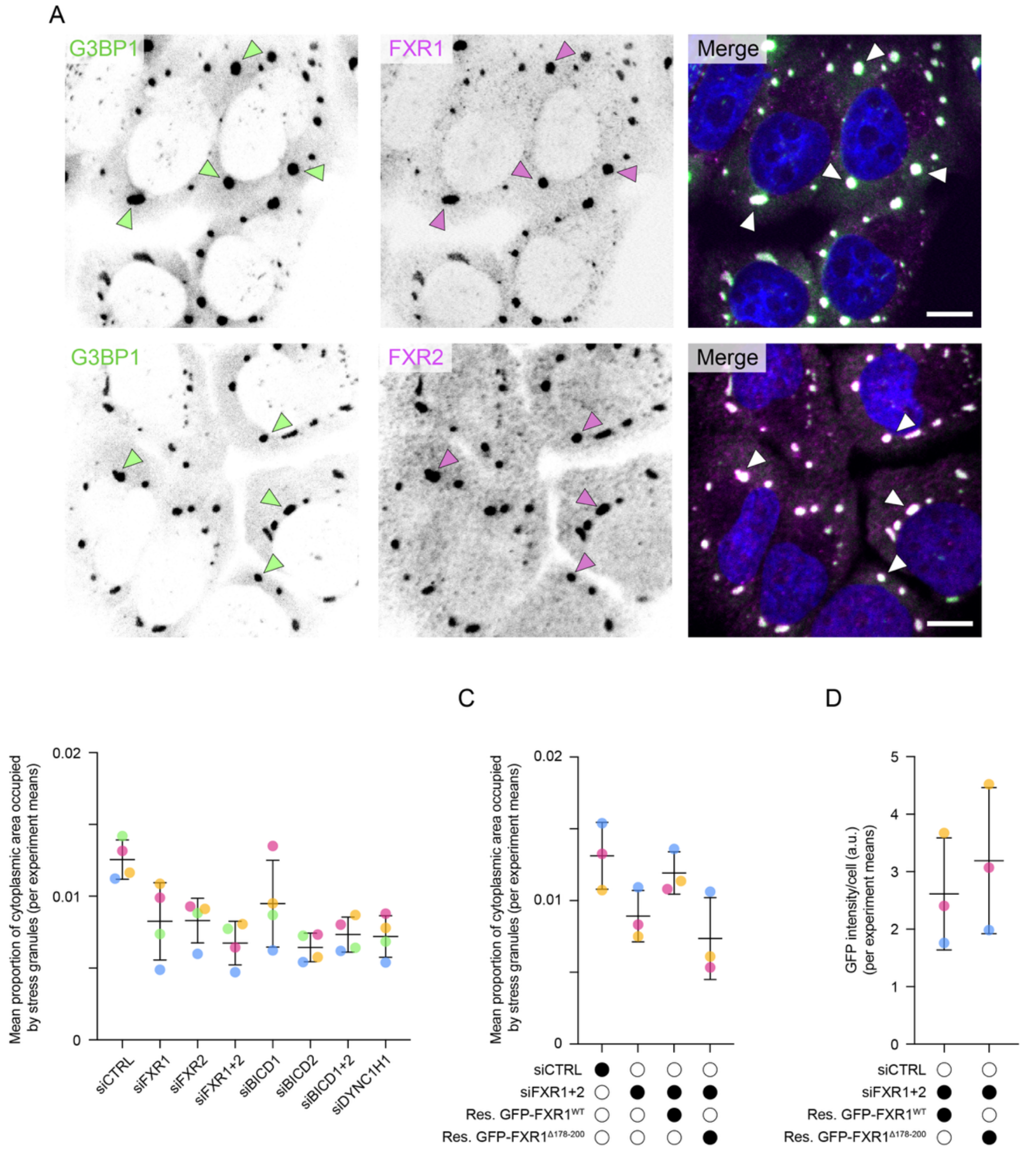


**Figure S9: Supplementary data on FXR proteins and stress granule assembly. (A)** Confocal images of immunostained U2OS cells showing enrichment of FXR1 and FXR2 in stress granules (arrowheads) that are formed by 1 h of arsenite treatment (300 µM) and detected with an antibody to endogenous G3BP1. Scale bar, 10 µm. **(B** and **C)** Mean values per experiment for proportion of cell area occupied by stress granules in the indicated conditions. Data correspond to those shown in Figure 6D (B) and Figure 6F (C). Within each panel, circles with the same color represent data from the same experiment. Note consistency of effects across almost all experiments. **(D)** Mean GFP intensity values per experiment for GFP-FXR$^{WT}$ and GFP-FXR$^{\Delta 178\text{-}200}$ transgenes in the experiments documented in Figure 6F and Figure S9C. Experiments are color-coded as in Figure S9C. In B–D, horizontal lines show means of the per experiment means and SD. Res., siRNA-resistant.

**Movie S1: Composite time series of stress granule assembly with siRNA treatments.** Image series from time-lapse microscopy showing stress granule formation (GFP-G3BP1) in U2OS cells treated with the indicated siRNA pools 72 h before addition of 300 μM arsenite for 1 h. The height of each image series corresponds to 54 μm.

**Table S1: Antibodies used in this study**

| | Target | Type | Supplier and Code | Dilution for IB | Dilution for IF |
|---|---|---|---|---|---|
| **Primary antibody** | β-actin | Mouse monoclonal IgM | Proteintech, 60008 | 1:2000 | - |
| | BICD1 | Rabbit polyclonal IgG | Abcam, ab170878 | 1:1000 | - |
| | BICD2 | Rabbit polyclonal IgG | Atlas Antibodies, HPA023013 | 1:1000 | 1:200 |
| | DYNC1I2 | Mouse monoclonal IgG | Sigma, MAB1618 | 1:1000 | - |
| | DYNC1H1 | Rabbit polyclonal IgG | Proteintech, 12345-1-AP | 1:500 | - |
| | eGFP | Chicken polyclonal IgY | Abcam, ab13970 | 1:5000 | - |
| | FXR1 | Rabbit polyclonal IgG | Atlas Antibodies, HPA018246 | 1:1000 | 1:200 |
| | FXR2 | Rabbit polyclonal IgG | Atlas Antibodies, HPA022997 | 1:1000 | 1:200 |
| | G3BP1 | Mouse monoclonal IgG | BD Transduction, 611126 | - | 1:200 |
| | MBP | Mouse monoclonal IgG | NEB, E8032 | 1:5000 | - |
| | DCTN1 | Mouse monoclonal IgG | BD Transduction Lab., 610474 | 1:2000 | - |
| | RAB6A | Rabbit polyclonal IgG | Atlas Antibodies, HPA059131 | 1:500 | - |
| | RPL8 | Rabbit monoclonal IgG | Abcam, ab169538 | 1:5000 | - |
| | RPS24 | Rabbit monoclonal IgG | Abcam, ab196652 | 1:5000 | - |
| **Secondary antibody** | Chicken | Goat IgY-HRP | Santa Cruz, sc-2428 | 1:5000 | - |
| | Mouse | Recombinant IgG-HRP | Santa Cruz, sc-516102 | 1:5000 | - |
| | Rabbit | Donkey IgG-HRP | GE Healthcare, NA934V | 1:5000 | - |
| | Chicken | Goat IgY, Alexa 488 | Invitrogen, A32931 | - | 1:1000 |
| | Mouse | Goat IgG, Alexa647 | Invitrogen, A21235 | - | 1:1000 |
| | Mouse | Goat IgG, Alexa488 | Invitrogen, A28175 | - | 1:1000 |
| | Mouse | Goat IgG, Alexa555 | Invitrogen, A28180 | - | 1:1000 |
| | Rabbit | Goat IgG, Alexa555 | Invitrogen, A21429 | - | 1:1000 |
| | Rabbit | Donkey IgG, Alexa488 | Invitrogen, A21206 | - | 1:1000 |
| | Rabbit | Goat IgG, Alexa647 | Invitrogen, A27040 | - | 1:1000 |

HRP, horseradish peroxidase; IB, immunoblotting; IF, immunofluorescence

**Table S2. RT-qPCR primers used in this study**

| Gene | Forward primer (5′–3′) | Reverse primer (5′–3′) |
|---|---|---|
| *G3BP1* | CGCGTAGGTTTGGACATATTTGACT | CCTGGTTCAGCAGTGTGTAA |
| *HSPA8* | TACCTTGGGAAGACTGTTACCAATG | GTCTAAGCCGTAAGCAATAGCAGCA |
| *TUBB* | AGATCGGTGCCAAGTTCTGG | GATCCACCAGGATGGCACGA |
| *XCR1* | GTGTAGATTCAGATGCTCTAAACGTCC | CACCAGGCAGTATAGGACAGTGGT |
